# A Chemical-Enhanced System for CRISPR-Based Nucleic Acid Detection

**DOI:** 10.1101/2021.03.28.437376

**Authors:** Zihan Li, Wenchang Zhao, Shixin Ma, Zexu Li, Yingjia Yao, Teng Fei

**Affiliations:** College of Life and Health Sciences, Northeastern University, Shenyang 110819, People’s Republic of China; Key Laboratory of Data Analytics and Optimization for Smart Industry (Northeastern University), Ministry of Education, Shenyang 110819, People’s Republic of China

**Keywords:** chemical, CRISPR, detection, diagnostics, SARS-CoV-2, COVID-19

## Abstract

The CRISPR-based nucleic acid detection systems such as SHERLOCK, DETECTR and HOLMES have shown great potential for point-of-care testing of viral pathogens, especially in the context of COVID-19 pandemic. Here we optimize several key parameters of reaction chemistry and develop a Chemical Enhanced CRISPR Detection system for nucleic acid (termed CECRID). For the Cas12a/Cas13a-based signal detection phase, we determine buffer conditions and substrate range for optimal detection performance. By comparing several chemical additives, we find that addition of L-proline can secure or enhance Cas12a/Cas13a detection capability. For isothermal amplification phase with typical LAMP and RPA methods, inclusion of L-proline can also enhance specific target amplification as determined by CRISPR detection. Using SARS-CoV-2 pseudovirus, we demonstrate CECRID has enhanced detection sensitivity over chemical additive-null method with either fluorescence or lateral flow strip readout. Thus, CECRID provides an improved detection power and system robustness towards practical application of CRISPR-based diagnostics.

## Introduction

Point-of-care testing (POCT) plays a pivotal role for infectious disease control by ensuring rapid and convenient diagnosis^1, 2^. The furious spreading of the novel coronavirus disease 2019 (COVID-19) caused by SARS-CoV-2 RNA virus has spurred a huge demand for reliable and field-deployable POCT solutions^3^. The recent advent of Clustered Regularly Interspaced Short Palindromic Repeats (CRISPR) technology not only revolutionizes genome editing field, but also represents a new paradigm for molecular diagnosis and POCT^4^. Several Cas nucleases such as Cas12a, Cas12b, Cas13a and Cas14 possess special collateral cleavage activity to cut nearby single stranded DNA or RNA molecules non-specifically, which is triggered by specific binding of guide RNA (gRNA, a chimeric RNA consisted of target-matching crRNA and trans-acting tracrRNA) along with CRISPR associated (Cas) protein complex to its cognate DNA or RNA target via precise base pairing. Taking advantage of this target-induced trans-cleavage activity as a signal amplifier, people have developed several CRISPR-based DNA or RNA detection methods including SHERLOCK (using Cas13a), DETECTR (Cas12a, Cas14), HOLMES (Cas12a) and CDetection (Cas12b)^5, 6, 7, 8, 9, 10, 11, 12^. Most of these methods include a target amplification phase to generate ample detection template from a trace amount of start material by varied isothermal amplification approaches such as loop-mediated isothermal amplification (LAMP) and recombinase polymerase amplification (RPA). In the following CRISPR detection phase, target-matching gRNA:Cas protein complex undergo conformational change to activate collateral cleavage onto nucleic acid reporter, thereby generating readout signal. The detection results are manifested by either portable fluorescence reader or lateral flow strips for POCT purpose. Inclusion of the target amplification phase can significantly increase the detection sensitivity, specificity and system robustness for CRISPR detection. The two phase reactions can be performed either separately or combined into one-tube reaction, although the latter may have compromised detection power due to constituent complexity and/or promiscuous reaction interference. Using these CRISPR-based methods, people have been trying to develop POCT solutions for monitoring microbial pathogens or genetic variants associated with diseases^13, 14^. During the COVID-19 pandemic, several CRISPR-based POCT methods for SARS-CoV-2 RNA detection were proposed^15, 16, 17, 18, 19, 20, 21, 22, 23, 24^ and a SHERLOCK-based COVID-19 test has been granted for an Emergency Use Authorization by The U.S. Food and Drug Administration recently^25^. However, these POCT solutions have not been widely applied yet, compared to the standard reverse transcription-polymerase chain reaction (RT-PCR) approach. Further development or improvement of these CRISPR detection methods is still necessary for broader and more practical applications in POCT settings.

Both CRISPR detection phase and isothermal target amplification phase are in essence composed of multiple enzyme-catalyzing processes. To elevate the detection power, people have been mainly focusing on selecting better effector components (e.g., Cas nuclease, gRNA, reporter, and polymerase), adjusting the stoichiometry of different constituents (e.g., Cas protein:gRNA ratio)^19, 25^ or modifying gRNA structure to augment trans-cleavage activity of Cas protein^18^. In addition to manipulating detection components, changing the reaction environment by either optimizing buffer condition (e.g., use Mn^2+^ instead of Mg^2+^) or adding chemical additives into reaction systems represent another directions to increase detection capability^25, 26^. Chemicals such as polyols, polymers, amino acid and their derivatives have been used as solvent additives to stabilize proteins and/or affect solvent properties, thereby influencing the kinetics of reaction chemistry^27^. Researchers have shown that chemical additives such as DMSO, glycerol and betaine can enhance the efficiency and/or specificity of some PCR or isothermal amplification reactions^28, 29, 30, 31, 32, 33^. However, systematic study is still required to examine whether these chemical additives may affect the detection power of CRISPR-based nucleic acid detection, in hope of searching the reaction enhancer for CRISPR diagnostics.

To determine the optimal reaction condition for Cas12a/Cas13a-mediated nucleic acid detection, here we examined several reaction parameters within the systems and evaluated several chemical additives for their effects on both target amplification and CRISPR detection phases. We found that L-proline can consistently enhance the reaction efficiency and specificity for both phases, and could serve as a practical additive for the reaction package of CRISPR-mediated nucleic acid detection. Applying this Chemical Enhanced CRISPR Detection system for nucleic acid (termed CECRID) in detecting SARS-CoV-2 pseudovirus demonstrated an improved detection power over additive-null methods, making CECRID a promising strategy towards further practical use.

## Results

### Optimization of key parameters for CRISPR detection systems

The DNA-targeting Cas12a and RNA-targeting Cas13a systems represent two major branches of the CRISPR machineries for nucleic acid detection (Fig. 1a). To compare and optimize these CRISPR detection systems, we set up three independent assays with purified AsCas12a, LbCas12a and LwaCas13a proteins (Methods and Supplementary Fig. 1). Fragments of ORF1ab, N and S genes from SARS-CoV-2 RNA genome were chosen as nucleic acid templates for detection. Target amplification regions by either LAMP or RPA method are denoted in Supplementary Fig. 2a. Different crRNAs for respective Cas effectors were designed and synthesized by in vitro transcription (Methods and Supplementary Fig. 2a). Fluorophore-labeled DNA reporter or RNA reporter were employed as signal readout for Cas12a and Cas13a systems, respectively (Fig. 1a).

**Fig. 1.**
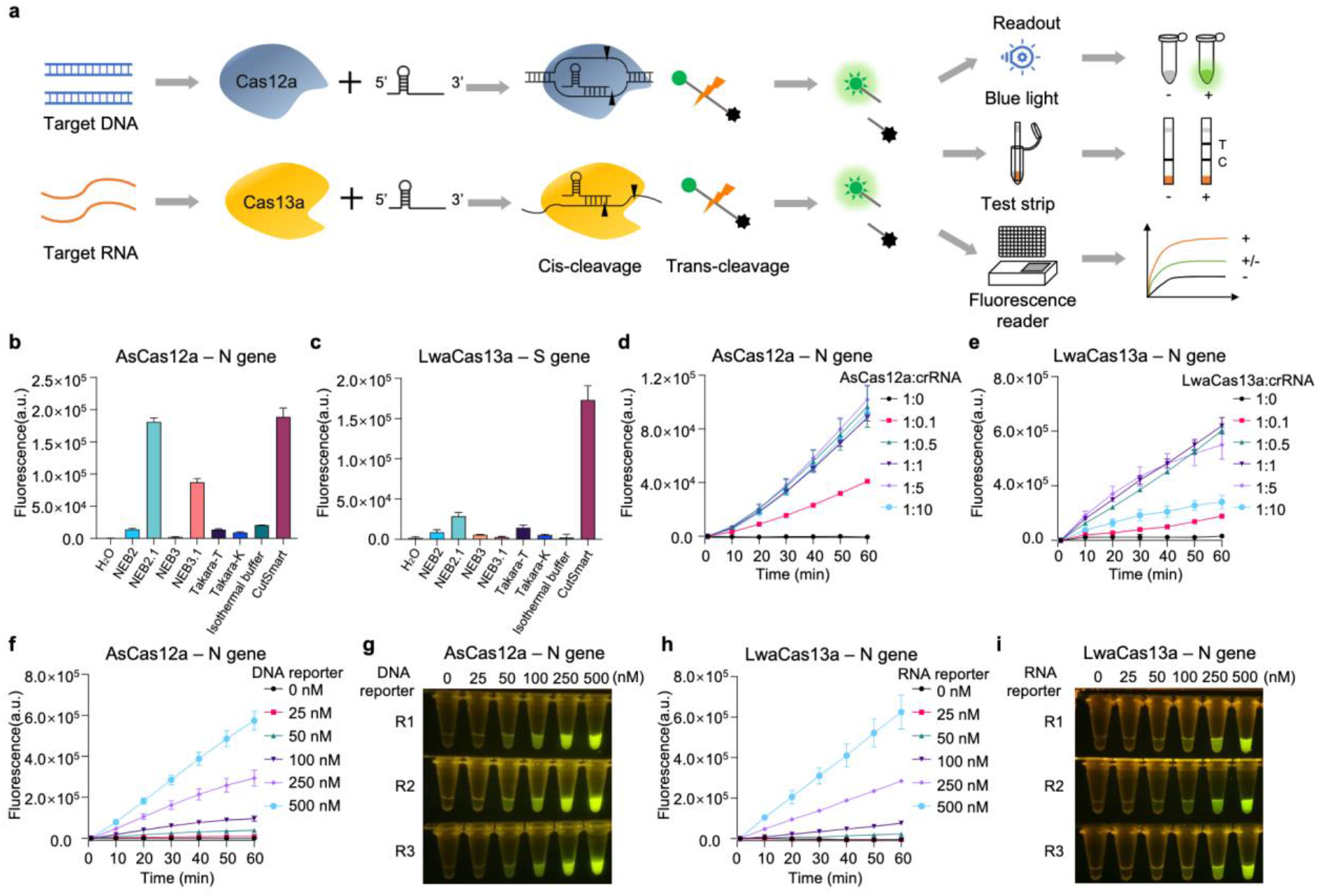
Optimization of key parameters for CRISPR detection systems. **a** Schematic description of CRISPR-Cas12a/Cas13a detection systems for nucleic acids. Specific binding of target DNA or RNA to Cas protein:crRNA complex triggers both the cis-and trans-cleavage activity of Cas nuclease. The latter activity is then employed to cleave an oligo-based reporter that can generate signals for detection. The representative ways to read out the signals include blue light illuminator, test strip and fluorescence reader. **b, c** Comparison of the indicated commercial reaction buffers for their effects on AsCas12a- and LwaCas13a-mediated trans-cleavage activity. AsCas12a-based CRISPR detection assays are set up with indicated buffers to detect a synthetic DNA template corresponding to SARS-CoV-2 N gene (b), LwaCas13a-based CRISPR detection assays are set up with indicated buffers to detect a synthetic RNA template corresponding to SARS-CoV-2 S gene (c). Error bars represent mean ± s.d. (n=3). a.u., arbitrary unit. **d, e** Evaluation of the indicated molecular ratio of Cas protein:crRNA for the effect on AsCas12a-based CRISPR detection using a synthetic DNA template corresponding to SARS-CoV-2 N gene (d), and LwaCas13a-based CRISPR detection using a synthetic RNA template corresponding to SARS-CoV-2 N gene (e). Error bars represent mean ± s. d. (n=3). a.u., arbitrary unit. **f-i** The effect of indicated amount of fluorescence reporter on CRISPR detection assays using either AsCas12a (f and g) or LwaCas13a (h and i). The results are shown by either real-time recording of fluorescence signal (f and h) or endpoint (60 min) visualization (g and i) under blue light illuminator. R1, R2 and R3 indicate three replicates. Error bars represent mean ± s.d. (n=3). a.u., arbitrary unit.

With synthetic ORF1ab and S DNA templates, we found that AsCas12a consistently has an enhanced detection sensitivity over LbCas12a in our hands as determined by the fluorescence signal (Supplementary Fig. 2b, c). We therefore mainly utilized AsCas12a for the following assays. To determine the optimal conditions for CRISPR-based detection, we evaluated several key parameters that affect the assay performance. Firstly, we compared different commercially available buffer systems for their effects on AsCas12a- or LwaCas13a-mediated detection with synthetic DNA or RNA templates. Interestingly, AsCas12a exhibited fair compatibility to several buffers (e.g., NEB2.1, NEB3.1 and CutSmart), whereas LwaCas13a seemed to be quite sensitive to buffer conditions (Fig. 1b, c). The commercial buffer CutSmart outperformed any other buffers tested for both Cas12a and Cas13a systems (Fig. 1b, c). Secondly, we determined the optimal range of Cas protein:crRNA ratio for the detection capability. AsCas12a displayed a saturated activity when the molecular ratio of Cas protein:crRNA sits between 1:0.5 ~ 1:10 (Fig. 1d). More crRNAs did not increase the fluorescence signal as AsCas12a amount is restricted (Fig. 1d). In contrast, LwaCas13a showed a narrower window of Cas protein:crRNA ratio (1:0.5 ~ 1:5) for maximal detection activity (Fig. 1e). Too much crRNA over LwaCas13a (at a ratio of 1:10) rather significantly decreased the fluorescence signal (Fig. 1e), suggesting the importance of appropriate stoichiometry of Cas effector and crRNA in Cas13a-mediated assays. Thirdly, we monitored the effect of DNA or RNA reporter amount on the detection signals. As expected, the fluorescence intensity was positively correlated with the amount of reporter within the tested range for both AsCas12a and LwaCas13a (Fig. 1f-i), These results suggest that the detection efficiency and signal strength could be elevated with increased amount of reporter on top of the fixed amount of other components.

### Dynamic detection range of target amount

Using the above optimized parameters, we determined the detection sensitivity of AsCas12a and LwaCas13a systems with pure DNA or RNA target templates. Both systems reached a significant signal detection threshold over the background when the target DNA or RNA had 10^9^ copies (~ 1.66× 10^-9^ mol/L in the reaction system) or more under indicated conditions in our hands (Fig. 2a, b). In practical scenarios of POCT, the start materials are often limited. To improve the detection efficiency and system robustness, people usually include an additional target amplification step by various isothermal amplification methods to generate ample target molecules for CRISPR detection. To determine the relationship between target abundance and CRISPR detection performance, we used differential amount of pure nucleic acid targets as CRISPR substrate and examined the corresponding signal strength. As the target substrate increased gradually, the detection signals from both AsCas12a and LwaCas13a initially correlated well with the target abundance, which is consistent with previous report^10^. However, the signals then started to drop after reaching the plateau as the targets continued to increase (Fig. 2c-f). It is surprising that too much redundant target substrates rather had an inhibitory effect, if not incremental or stable, on the detection system. We speculate that excess targets may competitively bind to crRNAs before the latter form complex with Cas proteins, thereby interfering the generation of productive target:gRNA:Cas protein complex. This effective target amount window is important for CRISPR detection systems to achieve optimal capacity and should be especially considered in assays conjugated with a target amplification phase.

**Fig. 2.**
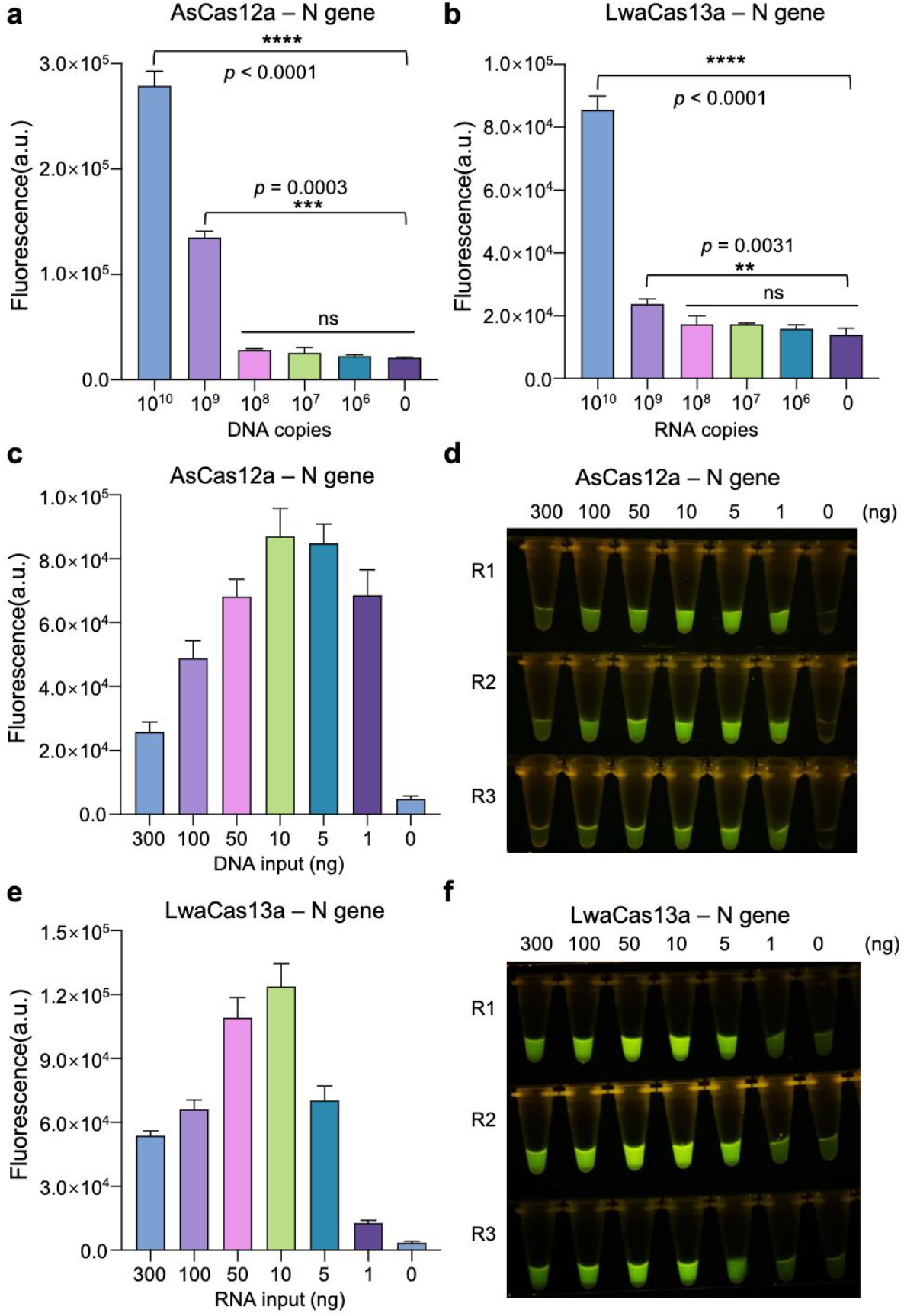
Dynamic detection range of target amount. **a, b** Evaluation of the detection sensitivity of AsCas12a (a) and LwaCas13a (b) systems with indicated amount of synthetic templates. Error bars represent mean ± s.d. (n=3). a.u., arbitrary unit. Unpaired two-tailed *t*-test, \*\**p* < 0.01, \*\*\**p* < 0.001, \*\*\*\**p* < 0.0001; ns means not significant. **c, d** AsCas12a-mediated detection power for indicated amount of pure targets using plasmid bearing SARS-CoV-2 N gene fragment. The fluorescence signals are shown by either real-time recording of fluorescence signal (c) or endpoint (60 min) visualization (d) under blue light illuminator. R1, R2 and R3 indicate three replicates. Error bars represent mean ± s. d. (n=3). a.u., arbitrary unit. **e, f** LwaCas13a-mediated detection power for indicated amount of pure targets using in vitro transcribed RNAs corresponding to SARS-CoV-2 N gene fragment. The fluorescence signals are shown by either real-time recording of fluorescence signal (c) or endpoint (60 min) visualization (d) under blue light illuminator. R1, R2 and R3 indicate three replicates. Error bars represent mean ± s.d. (n=3). a.u., arbitrary unit.

### Chemical additive enhancement for CRISPR detection phase

To further improve the efficiency of CRISPR detection systems, we tried to determine whether chemical additives may have some positive effects. Several widely used chemical additives were chosen and added into CRISPR detective reactions (Fig. 3a). For AsCas12a-mediated S gene DNA detection, addition of L-proline (0.2 M, 0.5 M and 1 M), DMSO (5%), and glycerol (1%, 5% and 10%) can greatly increase the detection efficiency (~ 1.6 to 6.68 fold) compared to control samples without chemical additive (Fig. 3b and Supplementary Fig. 3a). On the other hand, using N gene RNA template as substrate for LwaCas13a system, L-proline (0.2 M, 0.5 M and 1 M), betaine (0.5 M and 1 M) and DMSO (10%) displayed significant signal enhancement (~ 1.5 to 2.5 fold) over the control (Fig. 3c).

**Fig. 3.**
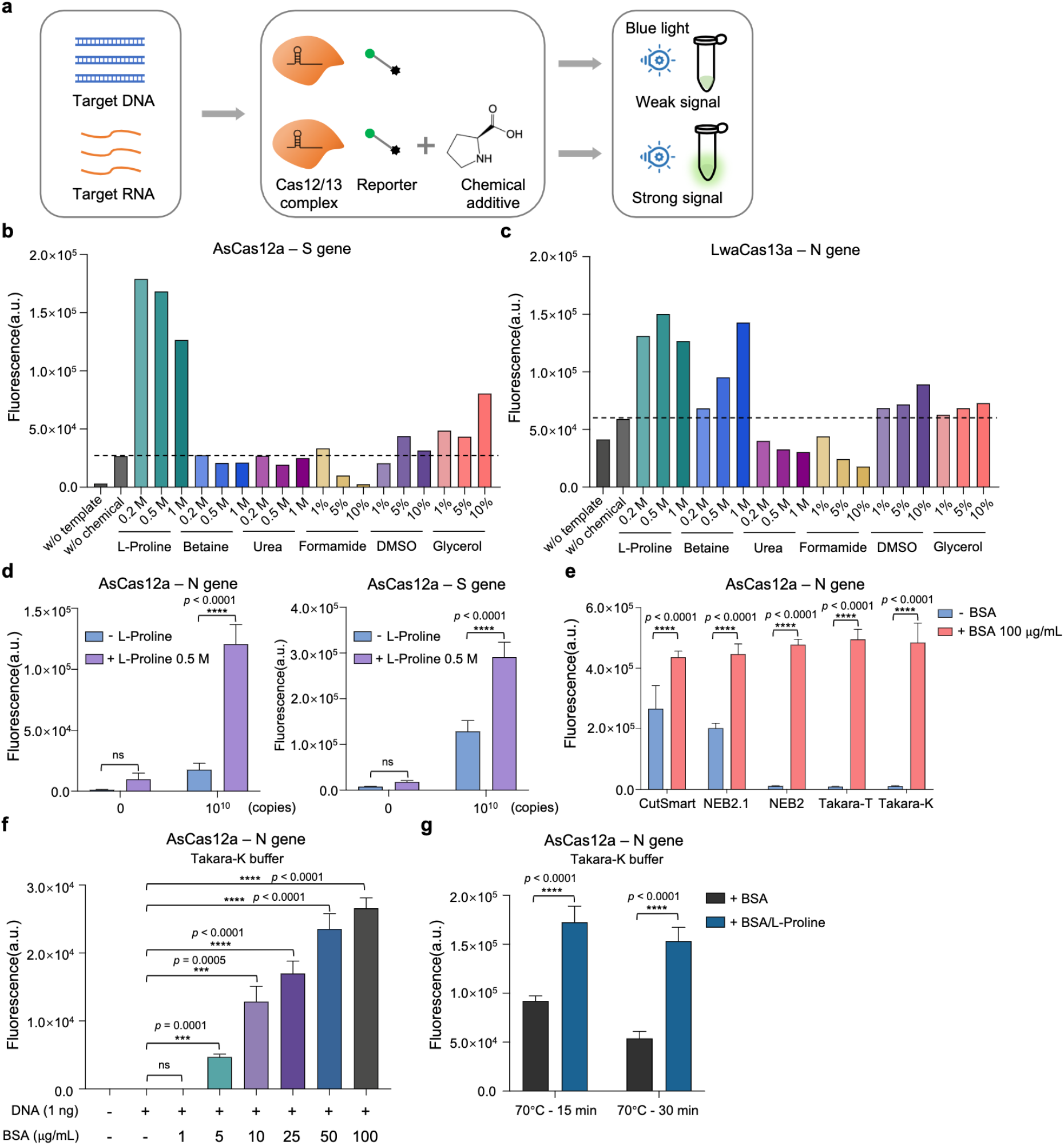
Effects of chemical additives on CRISPR detection phase. **a** Schematic diagram showing the workflow to evaluate chemical additive effect on CRISPR detection system. **b, c** Endpoint (60 min) recording of fluorescence detection signals for either AsCas12a (b) or LwaCas13a (c) system using synthetic S gene DNA template and N gene RNA template, respectively, with indicated amount of different chemical additives. The dotted line indicates the fluorescence baseline of CRISPR detection system without chemical addition using 10^10^ copies of SARS-CoV-2 nucleic acid targets. w/o template means no input in CRISPR detection system; w/o chemical means no chemical addition in the reaction of CRISPR system. **d** Comparison of fluorescent signals resulted from AsCas12a-based detection for SARS-CoV-2 N and S gene synthetic DNA fragments with or without 0.5 M L-proline addition. Error bars represent mean ± s.d. (n=3). a.u., arbitrary unit. Two-way ANOVA test, *\*\*\*\*p* < 0.0001; ns means not significant. **e** The effect of BSA addition on AsCas12a-based detection system under different reaction buffers using synthetic DNA fragment of SARS-CoV-2 N gene. Error bars represent mean ± s.d. (n=3). a.u., arbitrary unit. Unpaired two-tailed *t*-test, \*\*\*\**p* < 0.0001. **f** Comparison of different concentrations of BSA addition for the effects on AsCas12a-mediated detection of SARS-CoV-2 N gene synthetic DNA template using BSA-null Takara-K buffer. Error bars represent mean ± s.d. (n=3). a.u., arbitrary unit. Unpaired two-tailed *t*-test, \*\*\**p* < 0.001, \*\*\*\**p* < 0.0001; ns means not significant. **g** Evaluation of L-proline’s effect in protecting BSA from heat-induced denature and BSA’s capability to enhance AsCas12a-based CRISPR detection for SARS-CoV-2 N gene synthetic DNA template in Takara-K buffer. BSA along or BSA co-incubated with L-proline are treated at 70°C for 15 min or 30 min, before serving as additive in AsCas12a-mediated detection assays. Error bars represent mean ± s.d. (n=3). a.u., arbitrary unit. Two-way ANOVA test, \*\*\*\**p* < 0.0001.

Notably, L-proline, among all the tested chemicals, exhibited the most dramatic effect on both detection systems. We further verified the enhancement effect of 0.5 M L-proline on AsCas12a-based detection assays using both N and S gene DNA templates with replicates, suggestive of target- and crRNA-independence of L-proline’s enhancement effect (Fig. 3d). We postulate that L-proline may help to maintain or enhance the Cas effector activity as a protein stabilizer and refolding chaperone. To test this hypothesis, we prepared different types of AsCas12a protein with potentially differential basal activities (type 1: fresh protein from frozen stock; type 2: protein left at room temperature for 48 hours; and type 3: protein undergone multiple freeze-thaw cycles during 48 hours). As expected, the basal activities of AsCas12a protein in CutSmart buffer batch #1 declined gradually from type 1 to type 3 (Supplementary Fig. 3b). Interestingly, the addition of L-proline can significantly elevate the signal strength for all the three types of proteins in a dose-dependent manner (Supplementary Fig. 3b). Unexpectedly, when the similar tests were performed in CutSmart buffer batch #2 rather than buffer batch #1, the difference between basal activities of three types of AsCas12a are gone as type 2 and type 3 samples recovered their detection strength to the similar level of type 1 sample (Supplementary Fig. 3b, c). In addition, L-proline did not show enhancement effect in CutSmart buffer batch #2 (Supplementary Fig. 3c). These batch deviated results suggest that certain ingredients within CutSmart buffer #2 may retain the activity of AsCas12a even if after mild deterioration treatment, and L-proline’s effect on AsCas12a is probably masked by such endogenous signal recovery. In contrast, when AsCas12a protein was harshly denatured under 42°C heat stress, the decline of detection activities cannot be reversed back to normal range by the CutSmart buffer #2 (Supplementary Fig. 3d). However, addition of L-proline in detection mix retarded the heat-induced activity decline (Supplementary Fig. 3d), suggesting that L-proline may protect the activity loss of Cas nuclease against unfavorable stresses or conditions.

### BSA as an essential signal enhancer in CRISPR detection buffer

The significant batch effect of reaction buffers on the detection results and L-proline’s effect led us to interrogate the endogenous signal enhancers within the buffer. After scrutinizing and comparing buffer components between active and inactive buffers (Supplementary Table 1), we found that the presence or absence of bovine serum albumin (BSA) was highly correlated with the detection signal especially for AsCas12a (Fig. 1b, c). To test the potential function of BSA, we examined the detection activities of AsCas12a with N gene DNA template using different buffer systems while adding exogenous BSA. The tested buffers were chosen with similar salt and buffering constituents but mainly differed in the BSA content (BSA-inclusive buffers: CutSmart and NEB2.1; BSA-null buffers: NEB2, Takara-T and Takara-K) (Supplementary Table 1). As shown in Fig. 3e, the basal detection activity of BSA-inclusive buffers are significantly higher than BSA-null buffers, and addition of BSA into all the three BSA-null buffers drastically elevated their detection signals from the bottom to the saturated level. In contrast, adding L-proline alone into BSA-null buffers (Takara-T and Takara-K) cannot enhance the detection signal (Supplementary Fig. 3e). Furthermore, the enhancement effect of BSA is concentration-dependent, and the signal increase required the presence of target template which exclude the possibility that BSA’s addition had direct cleavage activity on the signal reporter (Fig. 3f and Supplementary Fig. 3f). Thus, BSA is a bona fide signal enhancer for CRISPR detection whereas L-proline may act as a safeguard in the buffer system to protect BSA or Cas protein from unfavorable stresses. Consistently, we did found that L-proline can protect BSA from heat stress and recovered the signal enhancement effect of BSA on CRISPR detection (Fig. 3g). Since BSA itself is a protein in essence and therefore requires more stringent manufacturing, transportation and storage conditions, the BSA-inclusive buffers such as CutSmart may display batch effect on CRISPR detection resulted from differential quality of endogenous BSA (Supplementary Fig. 3g, h). L-proline’s effect was more pronounced in buffers with low basal activity and probably deteriorated BSA’s function such as batch #3 and #6, and this phenomenon was consistently shown in both AsCas12a- and LwaCas13a-based assays (Supplementary Fig. 3g, h). The additional inclusion of L-proline in BSA-inclusive buffers may largely maintain the detection potential, antagonize the mild deterioration and represent the optimal reaction buffer system.

### Effects of chemical additives on target amplification phase

Pre-amplification of target molecules is often a requisite step before CRISPR detection phase to achieve the best detection sensitivity and specificity. LAMP and RPA are the most representative isothermal target amplification methods conjugated with CRISPR detection thus far (Fig. 4a). Using synthetic RNA template of SARS-CoV-2 N gene fragment, we performed LAMP-based target amplification. Total product yield accumulated quickly as reaction continued and apparent nonspecific product arose in non-template samples as determined by either ethidium bromide staining or fluorescent DNA binding dye quantification (Supplementary Fig. 4a, b). We then employed AsCas12a system to detect specific target signals, and found that LAMP-based two-step detection can significantly catch the signal from 10^3^ copies of template within 15 minutes of amplification (Supplementary Fig. 4c). Next, we tried to determine whether chemical additives could affect the efficiency and specificity of LAMP by inclusion of the tested additives within the reaction mix. As shown in Supplementary Fig. 4d, none of the tested chemicals can increase the gross yield of LAMP according to the signal kinetics captured by fluorescence reader using sequence-independent DNA-binding dye. Interestingly, when checking out the specific target amplification by CRISPR detection, only addition of L-proline (0.5 M and 1 M), among other tested chemicals, into LAMP reaction mix can significantly enhance the specific signal by ~4.5 fold (Fig. 4b, c). This enhancement effect was further reproduced by two independent LAMP-CRISPR two-step assays targeting SARS-CoV-2 N gene (Fig. 4d-k).

**Fig. 4.**
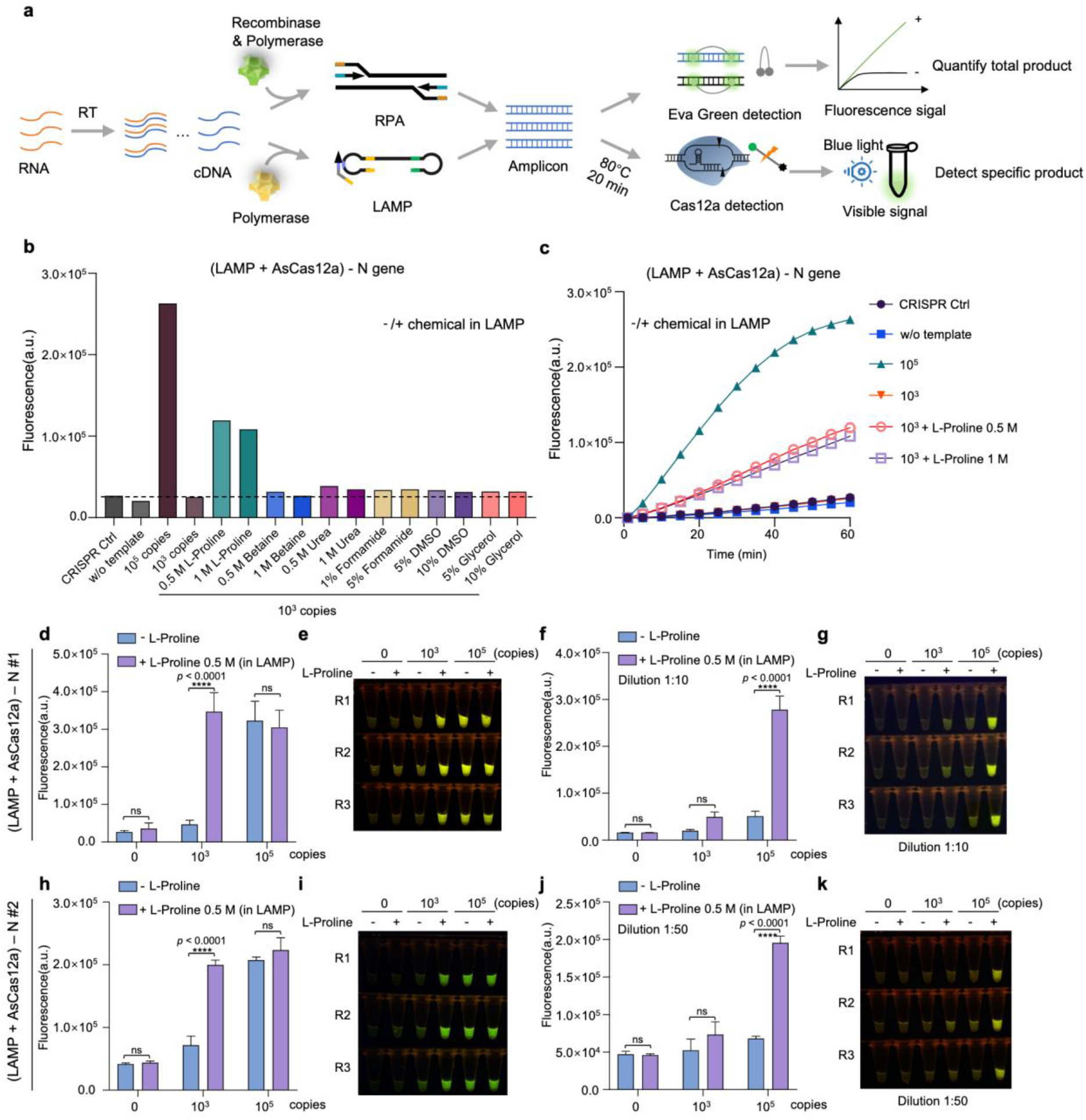
Effects of chemical additives on target amplification phase. **a** Schematic diagram showing the workflow to evaluate LAMP and RPA detection for target RNAs. Total amplification products are quantified by non-specific DNA-binding dye, and specific target amplification is determined by AsCas12a-mediated CRISPR detection. **b** Evaluation of different chemical additives for the effect on specific target amplification as determined by AsCas12a-based CRISPR detection using purified LAMP products (1:10 dilution) targeting SARS-CoV-2 N gene. Indicated chemicals are included in the LAMP reaction mix for a 15 min of reaction at 62°C. The LAMP products are purified and then subjected to CRISPR detection. Endpoint (60 min) recording of fluorescence detection signals are shown. The dotted line indicates the fluorescence baseline of CRISPR detection signal from the sample using 10^3^ copies of SARS-CoV-2 synthetic RNA template. CRISPR Ctrl indicates no nucleic acid input for AsCas12a detection system; w/o template means no input for LAMP reaction. **c** The fluorescence signal dynamics is shown for the selected samples in (b). **d-g** Comparison of L-proline addition in LAMP reaction phase for the effect on specific detection of SARS-CoV-2 N gene RNA template determined by AsCas12a detection system. The template and primer set for this LAMP reaction is quoted from Broughton et al and designated as N #1. Indicated amount of RNA template are used in LAMP system with or without 0.5 M L-proline for 30 min reaction at 62°C. After 20 min of 80°C heat inactivation, the LAMP products are purified and subjected to specific detection by AsCas12a system (d, e). Proper dilution (1:10) of purified DNA are performed to make sure the signals fall into effective detection range (f, g). The endpoint (60 min) fluorescence signal is shown by either bar plot (d, f) or direct visualization (e, g) under blue light illuminator. **h-k** Similar assays are performed as described in d-g, except that different template and primer set for LAMP reaction is used according to Joung et al.

We also performed similar analysis for another isothermal amplification method RPA (Fig. 4a). Both RPA and LAMP eventually produced significant background signals in empty control sample without target template as the run lasted to one hour, possibly due to the trade-off resulted from the easy and super-quick sparking principles of the methods (Supplementary Figs. 4a, b and 5a). However, these non-specific signals usually came late or weaker than target amplification wells, and target-derived specific signals still showed a time- and concentrationdependent increase by gRNA-mediated CRISPR detection (Supplementary Figs. 4c and 5b). Similar to LAMP, we did not find any chemical additive with significant enhancement effect on gross yield of RPA assay to amplify S gene RNA template (Supplementary Fig. 5c). On the other hand, for specific target amplification determined by two-phase RPA-CRISPR assays, we observed that several chemical additives such as L-proline (0.5 M and 1 M), betaine (0.5 M and 1 M), DMSO (5%) and glycerol (5%) displayed varied but significant boost of specific signals (Supplementary Fig. 5d-f). Again, L-proline showed the most comprehensive compatibility and effectiveness in reinforcing target amplification by either isothermal method in two-phase CRISPR detection assays. We also examined the effect of BSA addition on LAMP- and RPA-based assays using N gene RNA template. BSA addition into LAMP mix seemed to have little effect on specific target amplification, whereas in RPA assays BSA might have a beneficial role by promoting specific target amplification (Supplementary Fig. 6).

### Additional factors affecting signal-to-noise ratio of CRISPR detection

Given the high background nature of these fast isothermal amplification methods, it is critical to increase the signal-to-noise ratio for two-step CRISPR detection. In addition to chemical enhancement of the two phases, we also tried nested-RPA by introducing additional pair of RPA primers and performing two rounds of RPA amplification. Using ORF1ab gene RNA fragment as template, we observed ~8.9 fold increase of detection signal from nested-RPA-AsCas12a assay (Supplementary Fig. 7a). This nested primer approach may be less applicable for LAMP since there are already three pairs of primers with appreciable complexity within LAMP reaction system.

During our practice of two-phase CRISPR detection assay using RNA as template, we occasionally observed apparent background signals in both negative control and tested samples, which is not the case in assays with DNA template. Interestingly, this background signal can be removed if the input material from the pre-amplification step was heat-treated at 80°C for 20 minutes before proceeding to AsCas12a-mediated trans-cleavage assay (Supplementary Fig. 7b). Since DNA-targeting assay also contains the pre-amplification step but without such background signal, we reasoned that the key for this phenomenon might lie in the reverse transcription (RT) step which is a special difference between RNA- and DNA-targeting assays. We inferred that the input materials for CRISPR detection from RT-LAMP or RT-RPA step may contain some leftover of reverse transcriptase and dNTPs, which may cross-react with crRNAs in CRISPR detection mix to initiate promiscuous DNA production, thereby resulting in occasional background signals. To test this hypothesis, we added reverse transcriptase directly into LAMP-AsCas12a assay using N gene DNA template. The fluorescence signal was increased in reverse transcriptase-containing sample compared to control and 80°C heat-inactivation can effectively blunt this background increase, demonstrating that reverse transcriptase is the primary source of the background interference (Supplementary Fig. 7b, c, d). These results suggest that it may be advisable to take precautionary actions such as heat inactivation or DNA purification prior to CRISPR detection for reducing unwanted noise from reverse transcriptase in related assays.

### CECRID deployment for SARS-CoV-2 pseudovirus detection

With above optimized conditions and L-proline embedded in both target amplification phase and CRISPR detection phase, we established a Chemical Enhanced CRISPR Detection system for nucleic acid (termed CECRID) (Fig. 5a). To further validate the performance of CECRID in more practical settings, we packaged SARS-CoV-2 RNA fragments into VSVG-pseudotyped lentiviral particles to mimic live viruses for detection purpose. Quantified pseudoviral particles were directly added into viral transport media (VTM) to mimic samples collected from nasopharyngeal swab, and the viral RNA extraction was performed with commercial kit according to the standard procedures. The SARS-CoV-2 N gene viral RNA was detected by RT-LAMP-AsCas12a two-phase assays with two independent setup using different pseudoviruses/primers/crRNAs. Compared to standard assay without chemical additive, CECRID can significantly enhance the detection sensitivity of SARS-CoV-2 N gene RNA from pseudoviral particle input by either fluorescence reader or lateral flow strips (Fig. 5b-g). These results highlighted the potential of CECRID application in real POCT settings.

**Fig. 5.**
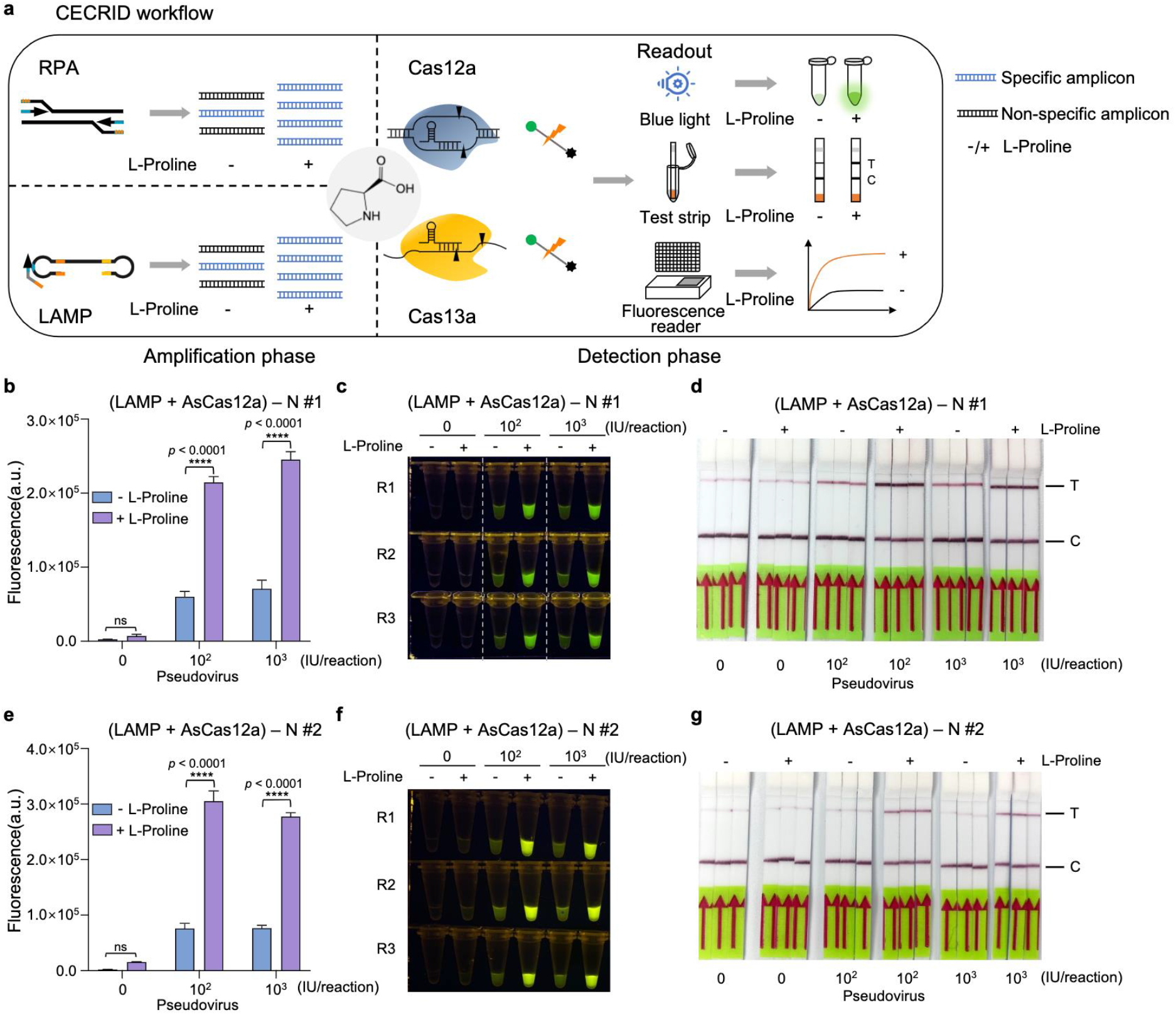
CECRID deployment for detection of SARS-CoV-2 pseudovirus. **a** Schematic diagram showing the principle and application workflow of CECRID systems. **b-d** Comparative analysis of SARS-CoV-2 N gene-bearing pseudovirus detection with CECRID and normal detection platforms. The template and primer set for this LAMP reaction is N#1. Various amount of pseudoviral particles (0, 10^2^ and 10^3^ IU, integration unit) in VTM are collected and viral RNA is purified by commercial RNA purification kit. RT-LAMP is performed with viral RNA at 62°C for 30 min followed by 20 min of 80°C inactivation. The purified amplification products (1:50 dilution) are then subjected to AsCas12a-based fluorescence and lateral flow strip detection. For CECRID, L-proline is added in both the LAMP phase and CRISPR detection phase. The endpoint (60 min) fluorescence signal is shown by either bar plot (b) or direct visualization (c) under blue light illuminator. An alternative readout is shown by the lateral flow strip using biotin-labeled reporter (d). The significant band in test line represents positive result. C: control line. T: test line. Error bars represent mean ± s.d. (n=3). a.u., arbitrary unit. Two-way ANOVA test, \*\*\*\**p* < 0.0001. **e-g** Similar assays are performed as described in b-d using an independent pseudovirus/template and primer set N #2. The purified amplification products (1:100 dilution) are then subjected to AsCas12a-based fluorescence and lateral flow strip detection. The endpoint (60 min) fluorescence signal is shown by either bar plot (e) or direct visualization (f) under blue light illuminator. An alternative readout is shown by the lateral flow strip using biotin-labeled reporter (g). Error bars represent mean ± s.d. (n=3). a.u., arbitrary unit. Two-way ANOVA test, \*\*\*\**p* < 0.0001.

## Discussion

In this study, we have explicitly revealed several basic features affecting the efficiency and sensitivity of different CRISPR detection systems. We further systematically evaluated several chemical additives for their effects on detection performance during either CRISPR detection phase or target amplification phase. By adding L-proline into the two-step CRISPR detection systems, we established an enhanced CRISPR detection toolkit named CECRID that exhibits improved system stability and detection power as evidenced using SARS-CoV-2 contrived gene template assays and pseudoviral particle testing. In addition, we also discussed some tips and procedures such as introducing nested amplification primers and reverse transcriptase inactivation towards improved CRISPR detection. These results will help to provide better CRISPR-based diagnostic solutions and expedite their practical applications for POCT purpose.

Chemical approaches have been adopted to modulate the activity of CRISPR/Cas system from several aspects: 1) chemical substance (e.g., 4-hydroxytamoxifen) can be used to achieve spatiotemporal control for Cas protein activity^34, 35^; 2) special chemical modifications are introduced into gRNA nucleotides for better stability, less off-target pairing or weaker immunogenicity^36^; 3) chemical engineering is applied to facilitate non-viral delivery of Cas protein:gRNA ribonucleoprotein complex into the cells^37^. All the above chemical engineering strategies are implemented to enhance the in vivo performance of CRISPR apparatus. For CRISPR-based in vitro application such as trans-cleavage-mediated nucleic acid detection, using chemical substance to boost CRISPR detection activity has not been thoroughly investigated. In contrast, several previous studies have tested the effects of chemical additives on in vitro target amplification by classic PCR. Some typical chemicals such as betaine, DMSO, BSA, dithiothreitol (DTT), and glycerol have been shown to improve the product specificity and/or yield during PCR amplification, especially for the difficult reactions with long or GC-rich template^28, 29, 30, 31, 32^. In addition, betaine was also reported to reduce non-specific product and enhance the amplification efficiency for isothermal RPA reaction^33^. However, betaine addition in LAMP reaction rather play an inhibitory role for target amplification, possibly due to the molecular barrier function of betaine to hinder intermolecular hybridization^38^. Joung et al recently established a one-pot SHERLOCK assay by combining LAMP-mediated target amplification and Cas12b-based CRISPR detection for SARS-CoV-2 viral detection^25^. They found that addition of either glycine or taurine into the one-pot reaction mix exhibited ~2-fold enhancement of the final detection signal^25^. However, these assays were only performed under one condition with specified primer/crRNA set, and the enzymatic constituents are quite complicated (including reverse transcriptase, DNA polymerase and Cas12b). Thus, it is still obscure to conclude whether the roles of these chemical additives are general for such type of reaction and on which step or component they exert the effect if any. Our work here systematically evaluated several commonly used chemical additives for their effects on either trans-cleavage-mediated CRISPR detection or LAMP/RPA-based isothermal target amplification, and identified a previously unrecognized chemical L-proline as a consistent chemical enhancer for these reactions.

General PCR enhancers such as betaine and DMSO are considered to play their roles by assisting double stranded DNA unwinding, lowering the melting temperature, reducing the formation of unwanted secondary structure between DNA strands, and facilitating primer annealing and extension^28, 29, 30, 31^. In contrast, BSA is not used as a typical PCR enhancer except for some difficult scenarios such as amplification from soil or plant samples that contain PCR inhibitor substance within the template^39, 40, 41^. BSA may bind to and neutralize the inhibitor as a blocking reagent, thereby promoting the amplification. BSA is usually regarded as a protein stabilizer by increasing the thermal stability and half-life of the enzymes during some typical molecular cloning-related reactions such as restriction enzyme digestion of DNA. In addition, human serum albumin or recombinant albumin was often included as a stabilizer in the formula for biological and medical reagents such as cytokine/hormone peptide and vaccine^42, 43^. Consistently, here we did not conclude a definite effect of BSA on isothermal amplification reactions despite some beneficial role for RPA assay with single specified primer/crRNA set. However, we did reveal a dramatic enhancement effect of BSA on CRISPR-based trans-cleavage reaction, suggesting that BSA should serve as a necessary component in CRISPR detection buffer. We posit that BSA may achieve this enhancement function through some or all of the following mechanisms: 1) stabilize Cas protein and other macromolecules in the reaction; 2) neutralize unknown inhibitory substance within the system especially from protein purification or nucleic acid preparation; 3) facilitate proper Cas protein refolding by modulating the solvent property; and 4) reduce the absorbance of reaction components on the test tube surface as a coating reagent.

L-proline is one of the most abundant molecules in cells and frequently used amino acids in natural proteins^44^. Proline residue plays an important role for protein folding via controlling the cis/trans isomerization of peptide bonds^45^. Pathogenic proline substitution is related to misfolding and aberrant aggregation of key protein presenilin 1 in Alzheimer’s disease^46^. In addition, proline-rich peptides were shown to possess immunomodulatory and neuroprotective properties against neurodegenerative diseases^47^. L-proline monomer can serve as natural osmoprotectant and cryoprotectant for cells exposed to osmotic and cold stresses possibly by preserving membrane integrity, stabilizing proteins and facilitating protein folding^44, 48^. Accordingly, L-proline was chosen as a stabilizer for liquid intravenous immunoglobulin (IVIG) products^49, 50^. Moreover, L-proline can serve as an efficient catalyst for several types of reactions in organic synthesis^51, 52, 53^. Here we found that L-proline, among other tested chemical additives, exhibits the most consistent effect in reinforcing the detection power during both the target amplification phase and CRISPR signal detection phase. Addition of L-proline significantly enhances the specific target amplification for LAMP and RPA reactions, which may be attributed to the multifaceted properties of L-proline such as stabilizing protein enzyme, assisting protein refolding, modulating the molecular interaction and creating favorable solvent environment. Interestingly, the effect of L-proline on CRISPR-mediated trans-cleavage depends on the buffer composition and even the buffer batch. Unlike BSA, which is a bona fide enhancer for CRISPR cleavage reaction, L-proline does not directly promote the CRISPR detection signal in less-favored buffers (e.g. BSA-null buffers) or certain batches of well-kept favored buffers (e.g. CutSmart buffer). Rather, L-proline displays an enhancer effect in the buffers when BSA is not in optimal conditions or the Cas protein enzyme is partially denatured. These results indicated that L-proline here may act as a system stabilizer to protect essential buffer component BSA and/or Cas enzyme from environmental stresses, thereby safeguarding their activities under suboptimal conditions. This notion is further supported by our data and previous reports showing that proline can prevent the aggregation of BSA and other proteins resulted from heat- or chemical-induced denaturation^54, 55^. Considering the complexity of the CRISPR reaction mix, the batch difference of reagent quality in applicable test kit is not only confined to the buffer system, but also expands to Cas enzymes, gRNAs and reporters throughout the whole manufacturing, transportation, storage and on-site testing processes. The inclusion of L-proline in the CRISPR detection mix provide an additional safety layer to secure the detection power and/or enhance the performance. More importantly, when CRISPR detection is conjugated with LAMP/RPA-based target amplification, L-proline as a system enhancer is compatible for both phases which simplifies the whole detection pipeline.

The enhancement effect of our CECRID platform have been reproduced with multiple Cas effectors, different target regions of templates, independent primer/crRNA sets and various vendors/batches of reagents, indicating the well generality and great applicability of the enhancement system. Despite our restricted access to real clinical samples, we corroborated the improved detection performance of CECRID for VTM-collected swab-like samples using SARS-CoV-2 pseudovirus. Taken together, our work not only uncovers several key parameters affecting the strength of nucleic acid detection, but also paves the way for developing robust reagent formula or test kit towards practical POCT application of CRISPR-based detection technology.

## Methods

### Constructs and reagents

The coding regions of LwaCas13a (*Leptotrichia wadeii* Cas13a)^56^, AsCas12a (*Acidaminococcus sp*. Cas12a)^57^ and LbCas12a (*Lachnospiraceae bacterium* Cas12a)^57^ were inserted into pET28a expression vector between BamHI and XhoI restriction enzyme sites for prokaryotic expression and purification of these proteins. N gene fragments (two different regions: #1 and #2) from SARS-CoV-2 viral genome were inserted into pHAGE-EF1α-puro vector between BamHI and KpnI restriction enzyme sites for SARS-CoV-2 pseudovirus detection. Oligonucleotides used in this study (Supplementary Table 2) were synthesized from HuaGene Biotech (Shanghai, China), Synbio Technologies (Suzhou, China) and GENEWIZ (Suzhou, China). Detailed information about reagents and instruments, including the commercial vendors and item numbers, is provided in Supplementary Table 3.

### Nucleic acid preparation

For preparation of DNA templates of SARS-CoV-2 ORF1ab, N and S genes, PCR amplification was performed by indicated primers with the forward primer containing an appended T7 promoter sequence using the template prepared through annealing of two synthetic oligonucleotides (Supplementary Table 2). For preparation of RNA templates, the in vitro transcription was performed with T7 promoter-inclusive DNA templates. Briefly, the in vitro transcription (IVT) system consisted of 4 μL 10x Transcription Buffer, 3.125 mM rNTPs, 50U T7 RNA Polymerase (Lucigen), 1 μL RNase Inhibitor and 50~100 ng DNA template, and the total reaction volume was 40 μL. After thorough mixing, the reaction system was incubated at 37°C for 1~2 hour (h). For preparation of crRNAs, the DNA oligonucleotide containing reverse complementary sequence of crRNA was annealed to an oligo (T7-F) with T7 promoter sequence to form a partial duplex DNA template for IVT. Then crRNAs were in vitro transcribed using the same IVT reaction system as described above. RNA templates and crRNAs were purified by RNA Clean & Concentrator-5 Kit (Zymo research), and quantified by the high sensitivity RNA Qubit fluorometer (Thermo Fisher).

### Cas protein expression and purification

Cas protein expression plasmids (pET28a-AsCas12a, pET28a-LbCas12a and pET28a-LwaCas13a) were transformed into *Escherichia coli*. Rosseta™ 2(DE3)pLySs competent cells. After transformation, cells were plated on kanamycin and chloramphenicol positive Luria-Bertani (LB) agar plate, and incubated for 16 h at 37°C. Pick up one colony from the plate, inoculate in 5 mL liquid LB medium supplemented with kanamycin and chloramphenicol, and put the starter culture on a shaker at 37°C overnight. 5 mL of starter culture was used to inoculate 1L of LB media supplemented with antibiotics and shaked at 37°C with 300 r.p.m. Cultures were allowed to grow until OD600 reached 0.4~0.6, and then cooled down for 30 minute (min) at 4°C. Add isopropyl β-D-thiogalactoside (IPTG) to a final concentration of 0.5 mM to induce protein expression for 14-16 h at 300 r.p.m. in a pre-chilled 20°C shaker. After induction, the cells were harvested by centrifugation (5000 g, 4°C, 10 min) for later purification and stored at −80°C.

Protein purification procedures were performed at 4°C. Cell pellet was resuspended in 15 mL of lysis buffer (20 mM Tris-HCl (pH 8.0), 500 mM NaCl, 1 mM DTT, 5% glycerol, 1 mM PMSF, 1 mg/mL lysozyme). The cell lysate was then sonicated by the sonicator with the following parameters: sonication (φ6, power 40%) for 1 second on and 2 seconds off with a total sonication time of 15 min. The sonicated sample was then centrifuged at 14000 r.p.m. for 10 min at 4°C and the supernatant was mixed with an equal volume of equilibration/wash buffer (50 mM sodium phosphate, 300 mM sodium chloride, 10 mM imidazole; pH 7.4). Since the recombinant Cas protein contains His-tag, HisPur™ Cobalt Resin (Thermo Fisher, #89964) was utilized to pull-down the protein. Washed protein extract was mixed with the prepared cobalt resin on end-over-end rotator for 30 min at 4°C followed by washing of the protein-bound cobalt resin twice in equilibration/wash buffer. Elute bound His tagged protein using elution buffer (50 mM sodium phosphate, 300 mM sodium chloride, 150 mM imidazole; pH 7.4) twice. Zeba™ Spin Desalting Columns (Thermo Fisher, #89890) was used to desalt protein, and ~2 mL protein could be collected after centrifuging at 850 g for 2 min. Mix the protein with 10 mL Storage Buffer (600 mM NaCl, 50 mM Tris-HCl (pH 7.5), 5% glycerol, 2 mM DTT), and transfer the mix into Amicon^®^ Ultra-15 Centrifugal Filter Devices (Millipore, #UFC905008) to concentrate the protein and exchange the storage buffer as well. The concentration of purified proteins was quantified by BCA protein assay kit (Meilunbio, #MA0082). SDS-PAGE analysis was performed with samples collected after different purification steps and the results were visualized by Coomassie Blue staining. Purified protein was stored at −80°C as 10 μL aliquots at a concentration of 2 mg/mL.

### CRISPR detection assay

CRISPR-LwaCas13a system was used for RNA detection. The standard LwaCas13a-based detection assay was performed at 37°C with 1x CutSmart buffer, 100 nM LwaCas13a protein, 100 nM crRNA, 250 nM RNA reporter and 1 μL of nucleic acid target in a 20 μL reaction system. For DNA detection system, CRISPR-AsCas12a and CRISPR-LbCas12a systems were applied. The standard reaction systems of Cas12a-based nucleic acid detection consisted of 1x CutSmart buffer, 50 nM AsCas12a protein or LbCas12a protein, 50 nM crRNA, 250 nM DNA reporter, and 1 μL of nucleic acid target or amplification products in a final volume of 20 μL at 37°C for 1 h. For the optimization of the detection systems, indicated amount of components and specified chemicals were added to the detection systems.

#### Fluorescence readout

The CRISPR-LwaCas13a and CRISPR-AsCas12a (LbCas12a) fluorescence detection assays were performed by using fluorophore-quencher (FQ) reporters involving a short single-stranded (ss) RNA or DNA oligonucleotide, respectively. Both of the ssRNA and ssDNA FQ reporters were composed of a 6-FAM fluorophore on 5 terminal and a BHQ1 quencher on 3 terminal (ssRNA FQ reporter: 5’ -/6-FAM/UUUUUU/BHQ1/-3’; ssDNA FQ reporter: 5’ -/6-FAM/TTATT/BHQ1/-3’). For LwaCas13a-based assay, the detection system consisted of 100 nM LwaCas13a, 100 nM crRNA, 250 mM ssRNA fluorescent reporter, 1x CutSmart Buffer and 1 μL nucleic acid target in a 20 μL reaction system. For AsCas12a- and LbCas12a-based assays, the detection system contained 50 nM AsCas12a or LbCas12a protein, 50 nM crRNA, 250 nM ssDNA fluorescence reporter, 1x CutSmart Buffer and 1 μL DNA target or amplification products. Fluorescence signal was dynamically measured by QuantStudio™ 5 Real-Time PCR System (Thermo Fisher). Background-subtracted signals for each monitoring points were further normalized by subtraction of its initial value to make comparison between different conditions (arbitrary unit, a.u.) for the analysis. Visual detection was accomplished by imaging the tubes through E-Gel™ Safe Imager™ Real-Time Transilluminator (Thermo Fisher).

#### Lateral flow readout

For lateral flow strip assays, the AsCas12a-based detection system was assembled as described above except that the fluorescence reporter was replaced with the biotin-labeled reporter. Lateral flow cleavage reporter (5’ -/6-FAM/TTATTATT/Biotin/-3’) was added to the reaction at a final concentration of 250 nM in 20 μL reaction volume along with the 1 μL RT(reverse transcription)-LAMP product, and incubate the reaction at 37°C for 1 h. After completion of the incubation, add 80 μL HybriDetect Assay Buffer into the reaction tube. A lateral flow strip (Milenia Biotec GmbH, # MGHD 1) was placed to the reaction tube and incubated for approximately 3~5 min, and the result was visualized directly. The negative result is indicated by one primary band at the control line (C), and significant band in test line (T) represents positive result.

### LAMP assay

For detection of SARS-CoV-2 RNA, RT-LAMP reactions were performed with caution on a dedicated clean bench using filter tips to prevent sample contamination. One-step RT-LAMP mix was assembled with 1 μL 10x isothermal amplification buffer, 1.4 mM of dNTPs, 6.5 mM MgSO4, 3.2 U Bst 2.0 (NEB, #M0538S), 3 U WarmStart RTx Reverse Transcriptase (NEB, #M0380S), 0.2 μM F3/B3 primers, 1.6 μM FIP/BIP primers, 0.8 μM LF/LB primers, and 1 μL of RNA template in a 10 μL volume, and then incubated at 62°C for 15~30 min. Primers for LAMP are listed in Supplementary Table S2. The heat inactivation (80°C for 20 min) of LAMP product was performed to reduce the background signal before proceeding to CRISPR detection assays. Quantitative real-time PCR was conducted using non-specific DNA-binding dye EvaGreen (Biotium, #31000) to quantitate the total amount of amplification products. EvaGreen-derived fluorescence signal was normalized by subtraction of initial value to make comparison between different conditions. For evaluation of the target specificity of LAMP reactions, amplification products were purified by PCR Purification Kit (Thermo Fisher, #K0702), and followed by fluorescence detection with AsCas12a system. The purified products might be diluted to an appropriate concentration to make sure the detection signal fell into the effective detection range.

### RPA assay

RPA assays were set up using commercial RPA kit (TwistDx). One step RT-RPA reaction system was composed of 9 μL of RPA solution (primer-free rehydration buffer), 224 mM of MgOAc, 40 U of ProtoScript II Reverse Transcriptase (NEB, #M0368S), 0.5 μM of forward primer, 0.5 μM of reverse primer and add UltraPure water to 16 μL. Primers for RPA are listed in Supplementary Table S2. The mixture was incubated at 40°C for 30 min, followed by heat inactivation for 20 min at 80°C. Total amount of amplification product was quantified by quantitative real-time PCR using EvaGreen dye, and specific target amplification was determined by AsCas12a-based CRISPR detection after purification using PCR Purification Kit.

### Pseudovirus production and detection

HEK293FT cells were employed to pack the SARS-CoV-2 pseudoviral particles. Cells were cultured with DMEM medium supplemented with 10% fetal bovine serum. DNA Transfection was performed in 6-well plates by Lipofectamine™ 2000 Transfection Reagent with a mix of 1.5 μg pHAGE-EF1α-puro plasmid carrying SARS-CoV-2 N gene fragment (#1 and #2), 0.75 μg pCMVR8.74, and 0.5 μg pMD2.G. The viral particle-containing supernatant was harvested at 48 h post transfection and centrifuged at 3000 r.p.m for 5 min to remove the cell debris. Aliquot and store the virus supernatant at −80°C before use. For virus titration, a commercial Lentivirus Titer Kit was used (Abm, #LV900). Briefly, 1 μL virus supernatant was lysed in 9μL Virus Lysis Buffer for 3 min at room temperature. Quantitative Reverse Transcription PCR (qRT-PCR) was performed with specified primer set to quantify the viral particles. The titer of virus can be calculated from on-line lentiviral titer calculator which is provided by Abm at http://www.abmgood.com/High-Titer-Lentivirus-Calculation.html. To simulate the actual application, viral particles were firstly transferred into the Viral Transport Media (VTM) of Sample Collection Kit (BEAVER, #43903) to a final volume of 1 mL as a mimic of nasopharyngeal swab collected sample. Take out 100 μL solution for RNA extraction by UNIQ-10 Column Trizol Total RNA Isolation kit (Sangon, #B511321-0100), and elute the RNA by 50 μL Ultrapure H2O. RT-LAMP was performed with 1 μL extracted RNA as template followed by AsCas12a-based detection in the absence or presence of L-proline.

### Statistical analysis

Statistic significances were calculated by GraphPad Prism 8.4.0 and all the data were shown as mean ±s.d. The two-way ANOVA with Sidak’s multiple comparisons test was used to compare differences between groups. Statistical significance was determined by an unpaired two-tailed t-test. Asterisks indicate \*\**p* < 0.01, \*\*\**p* < 0.001, \*\*\*\**p* < 0.0001.

## Acknowledgements

We thank Drs. Tengfei Xiao, Yue Feng and Ren Sheng for sharing reagents. This work was supported by the National Natural Science Foundation of China (31871344; 32071441), the Fundamental Research Funds for the Central Universities (N182005005; N2020001), the 111 Project (B16009), and LiaoNing Revitalization Talents Program (XLYC1807212).

## Author Contributions

Zihan.L. and T.F. conceived and designed the research. Zihan.L, W.Z., and S.M. performed the research. All the authors analyzed the data. Zihan.L, W.Z., S.M. and T.F. wrote the manuscript with the help of all the other authors. T.F. supervised the study.

## Supplementary Information

Supplementary Fig. 1 Expression and purification of Cas protein.

Supplementary Fig. 2 Schematic of SARS-CoV-2 genome and comparative analysis of trans-cleavage activity between two Cas12a ortholog proteins.

Supplementary Fig. 3 Effects of chemical additives, BSA and buffer batch on CRISPR detection.

Supplementary Fig. 4 Evaluation of LAMP and chemical additives for target detection.

Supplementary Fig. 5 Evaluation of RPA and chemical additives for target detection.

Supplementary Fig. 6 The effect of BSA on target amplification of LAMP and RPA.

Supplementary Fig. 7 Performance of RT-Nested RPA and background removal for CRISPR detection.

Supplementary Table 1 The constitutions of different reaction buffers.

Supplementary Table 2 Oligonucleotides used in this study.

Supplementary Table 3 Reagents and instruments used in this study.

**Supplementary Fig. 1.**
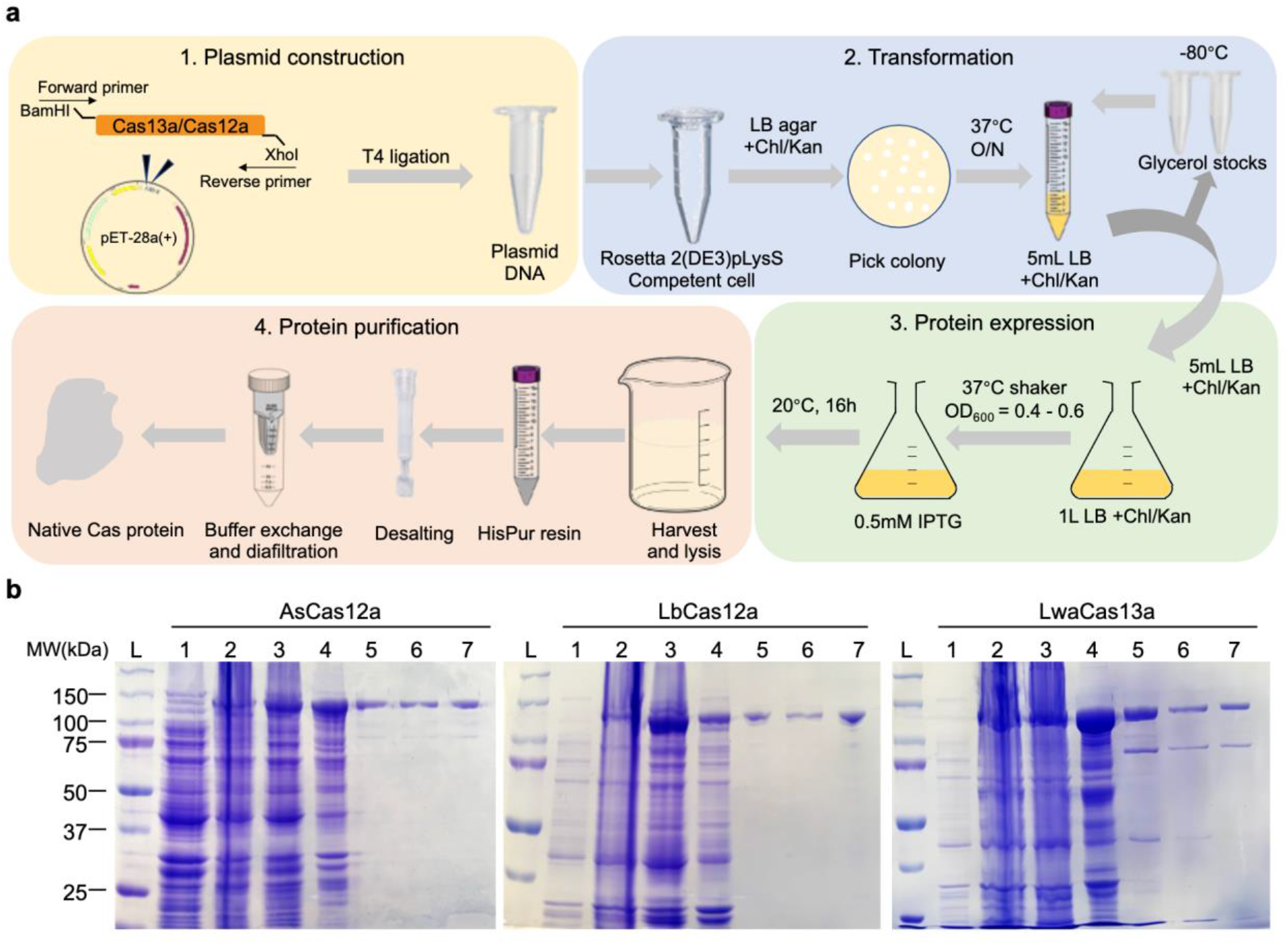
Expression and purification of Cas protein. **a** Schematic workflow of protein expression and purification of Cas nucleases. **b** Different protein fractions collected during protein purification are analyzed by SDS-PAGE and followed by Coomassie Blue staining. L: ladder; 1: non-induced; 2: cell lysate; 3: supernatant of lysate; 4: pellet of lysate; 5: eluted fraction post HisPur Cobalt Resin; 6: fraction post desalting; 7: final purified product post concentrating and buffer exchange.

**Supplementary Fig. 2.**
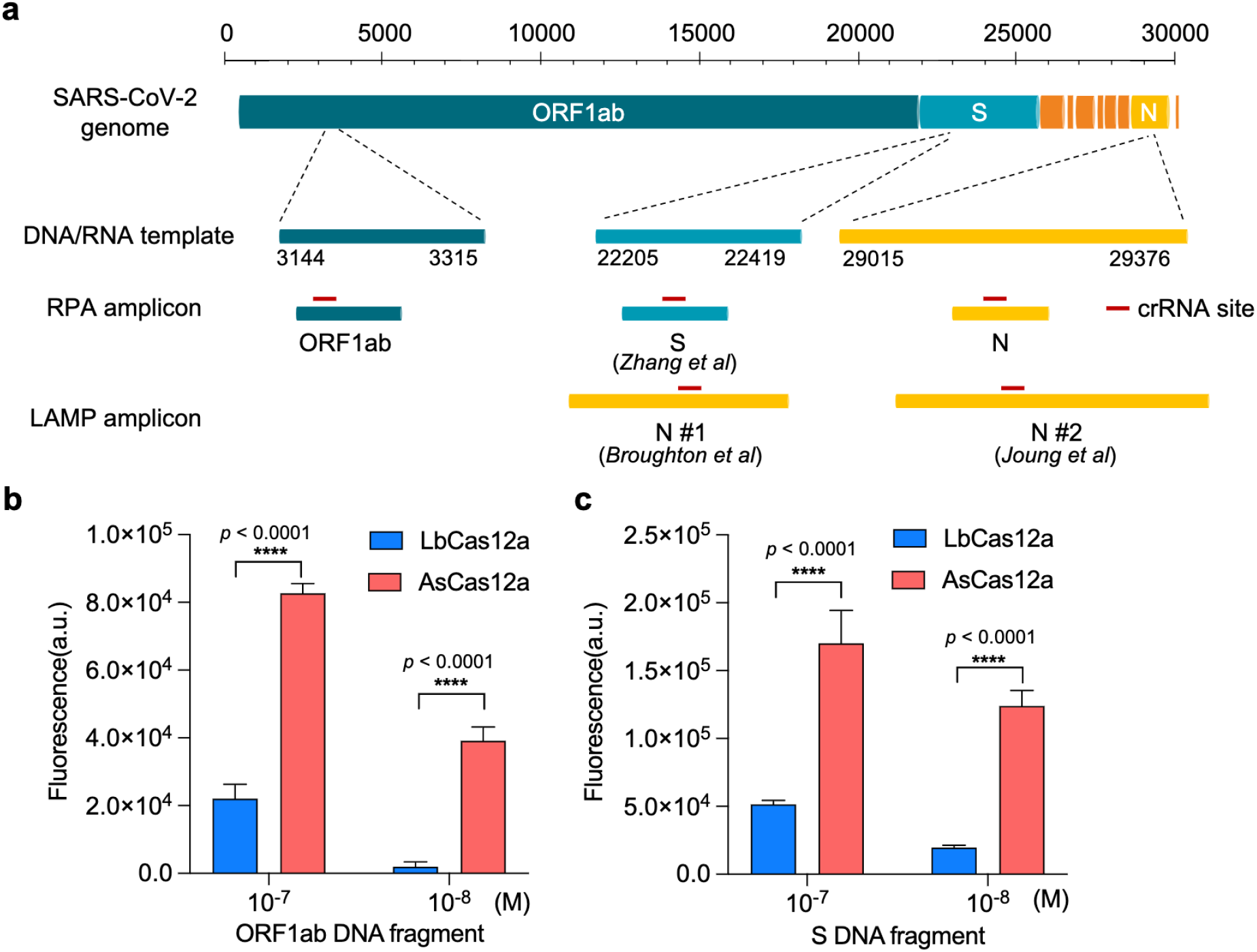
Schematic of SARS-CoV-2 genome and comparative analysis of trans-cleavage activity between two Cas12a ortholog proteins. **a** Schematic diagram showing the genomic architecture of SARS-CoV-2 virus. The positions and regions for DNA and RNA templates, RPA and LAMP amplicons, and crRNA-targeting sites used in this study are donated. ORF1ab: open reading frame 1ab; S: spike protein; N: nucleocapsid protein. **b** Comparison of two Cas12 ortholog proteins (LbCas12a and AsCas12a) for trans-cleavage-mediated detection of synthetic SARS-CoV-2 ORF1ab gene DNA template. Endpoint (60 min) recording of fluorescence detection signals are shown. Error bars represent mean ±s. d. (n=3). a.u., arbitrary unit. Two-way ANOVA test, \*\*\*\**p* < 0.0001. **c** Comparison of two Cas12 ortholog proteins (LbCas12a and AsCas12a) for trans-cleavage-mediated detection of synthetic SARS-CoV-2 S gene DNA template. Endpoint (60 min) recording of fluorescence detection signals are shown. Error bars represent mean ±s. d. (n=3). a.u., arbitrary unit. Two-way ANOVA test, \*\*\*\**p* < 0.0001.

**Supplementary Fig. 3.**
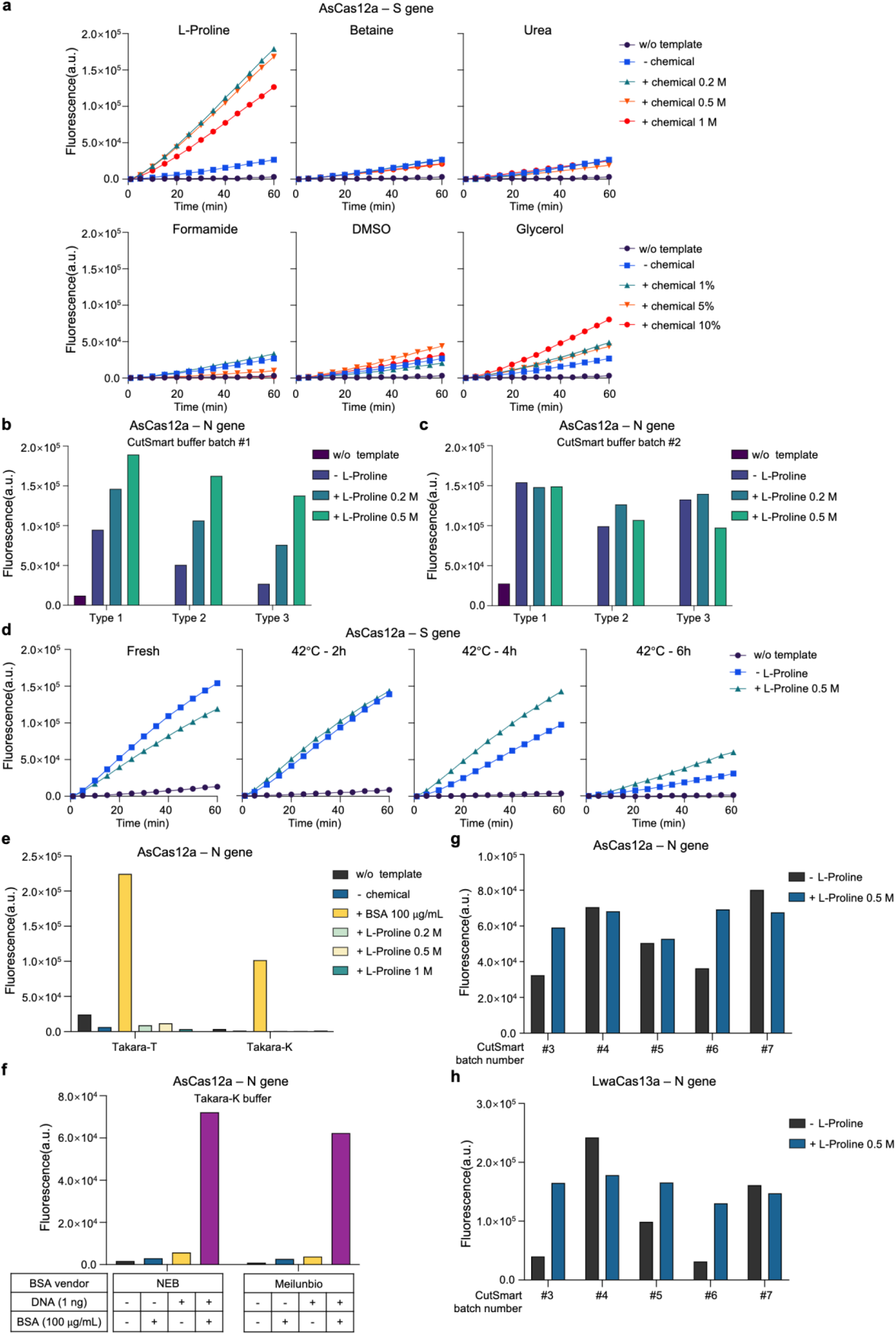
Effects of chemical additives, BSA and buffer batch on CRISPR detection. **a** Fluorescence signal kinetics of AsCas12-mediated detection for SARS-CoV-2 S gene synthetic DNA (10^10^ copies) with indicated chemical additives in the reaction mix. w/o template means no input in CRISPR detection system; differential amount of chemicals are added in the reaction mix: no chemical (-), with indicated amount of chemical (+). a.u., arbitrary unit. **b** Effects of L-proline on AsCas12-mediated SARS-CoV-2 N gene DNA template detection using different types of Cas protein representing differential states of denaturing in batch #1 CutSmart reaction buffer. L-proline is added in the CRISPR detection mix. Three types of AsCas12a protein are used in this assay - type 1: fresh protein from frozen stock; type 2: protein left at room temperature for 48 hours; and type 3: protein undergone multiple freeze-thaw cycles during 48 hours. Endpoint (60 min) recording of fluorescence detection signals are shown. w/o template means no input in CRISPR detection system; differential amount of L-proline are added in the reaction mix: no L-proline (-), with indicated amount of L-proline (+). a.u., arbitrary unit. **c** Batch effect of reaction buffers on AsCas12a-based detection and L-proline’s enhancement. The same samples used in **b** are gone through the similar assays only except that the reaction buffer changes to batch #2. a.u., arbitrary unit. **d** Effects of L-proline on AsCas12a-based detection for SARS-CoV-2 S gene DNA template with heat-denatured AsCas12a proteins. AsCas12a protein is pre-heated for 2, 4 and 6 hours in 42°C, and then is examined for their capability on CRISPR detection in the absence (-) or presence (+) of 0.5 M L-proline in the detection mix. Fluorescence signal kinetics is shown. a.u., arbitrary unit. **e** Effects of BSA and L-proline addition on AsCas12-mediated SARS-CoV-2 N gene DNA template detection using Takara-T and Takara-K buffers. Endpoint (60 min) recording of fluorescence detection signals is shown. w/o template means no input in CRISPR detection system; differential amount of chemicals are added in the reaction mix: no chemical (-), with indicated amount of chemical (+). a.u., arbitrary unit. **f** Comparison of different sources of BSA on AsCas12-mediated SARS-CoV-2 N gene DNA template detection in Takara-K buffer. Different vendors of BSA are used. Endpoint (60 min) recording of fluorescence detection signals are shown. a.u., arbitrary unit. **g, h** Evaluation of multiple batches (#3 - #7) of CutSmart buffer for the effect on AsCas12a (g) and LwaCas13a (h) detection capability without (-) or with (+) 0.5 M L-proline addition using SARS-CoV-2 N gene DNA (g) or RNA (h) templates. Endpoint (60 min) recording of fluorescence detection signals are shown. a.u., arbitrary unit.

**Supplementary Fig. 4.**
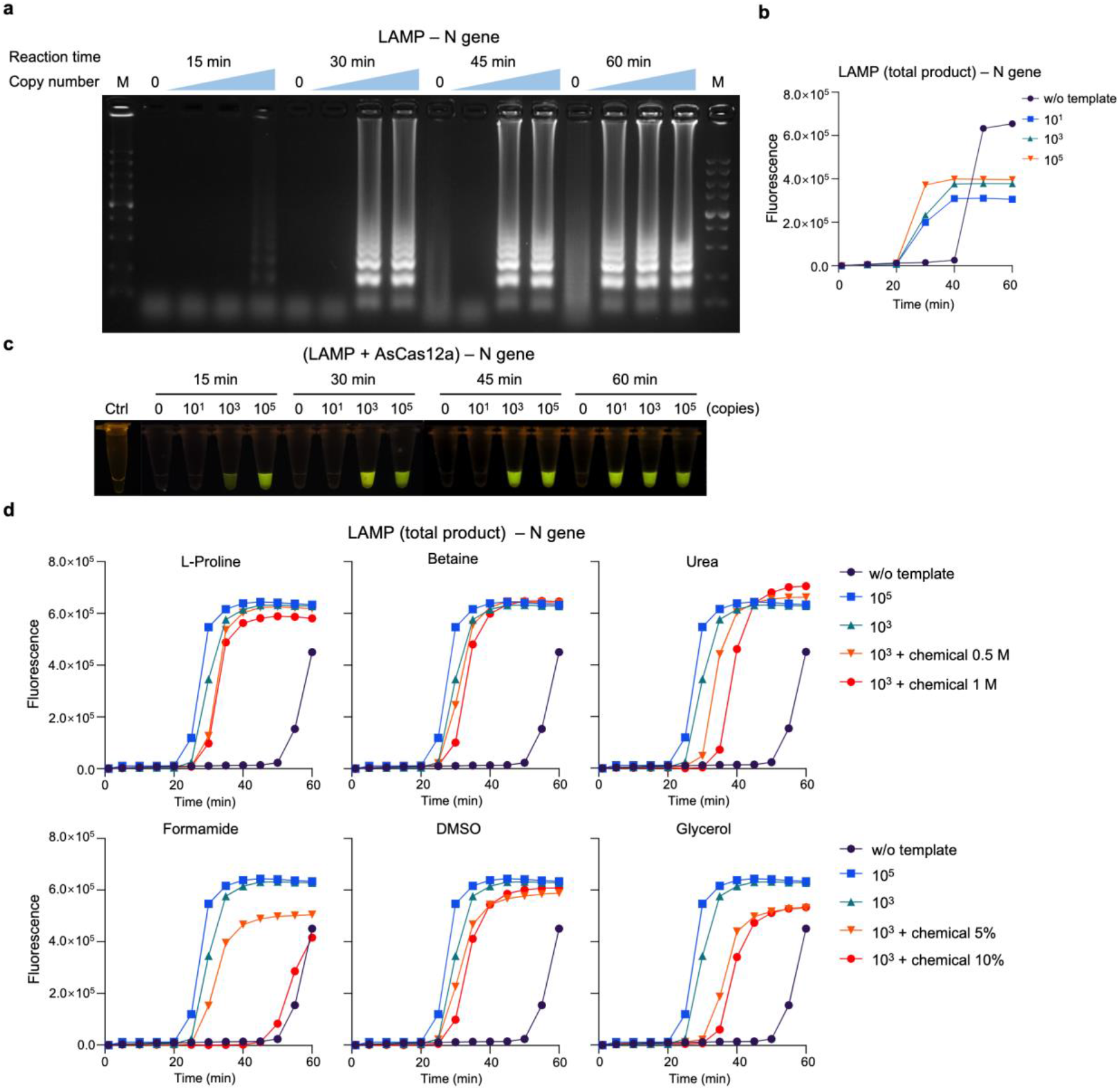
Evaluation of LAMP and chemical additives for target detection. **a** Agarose gel electrophoresis of the RT-LAMP products using different amount of SARS-CoV-2 N gene RNA template with different time duration of reactions. LAMP is performed at 62°C for 15, 30, 45 and 60 min. RNA template copy number (left to right): 0, 10^1^, 10^3^, 10^5^ copies. **b** Dynamic monitoring of total amplification product of RT-LAMP by nonspecific DNA-binding dye EvaGreen using varied amount of SARS-CoV-2 N gene RNA template. **c** Specific target amplification is determined using purified LAMP products by AsCas12a-based detection with fluorescence signal directly visualized by blue light illuminator. The samples are the same as used in (a). **d** Evaluation of chemical additives for the effects on total products of RT-LAMP using SARS-CoV-2 N gene RNA template (10^3^ and 10^5^ copies). Indicated chemicals (+) are added in the LAMP reaction mix and fluorescence kinetics is shown as determined by EvaGreen dye. w/o template means no template input in LAMP reaction.

**Supplementary Fig. 5.**
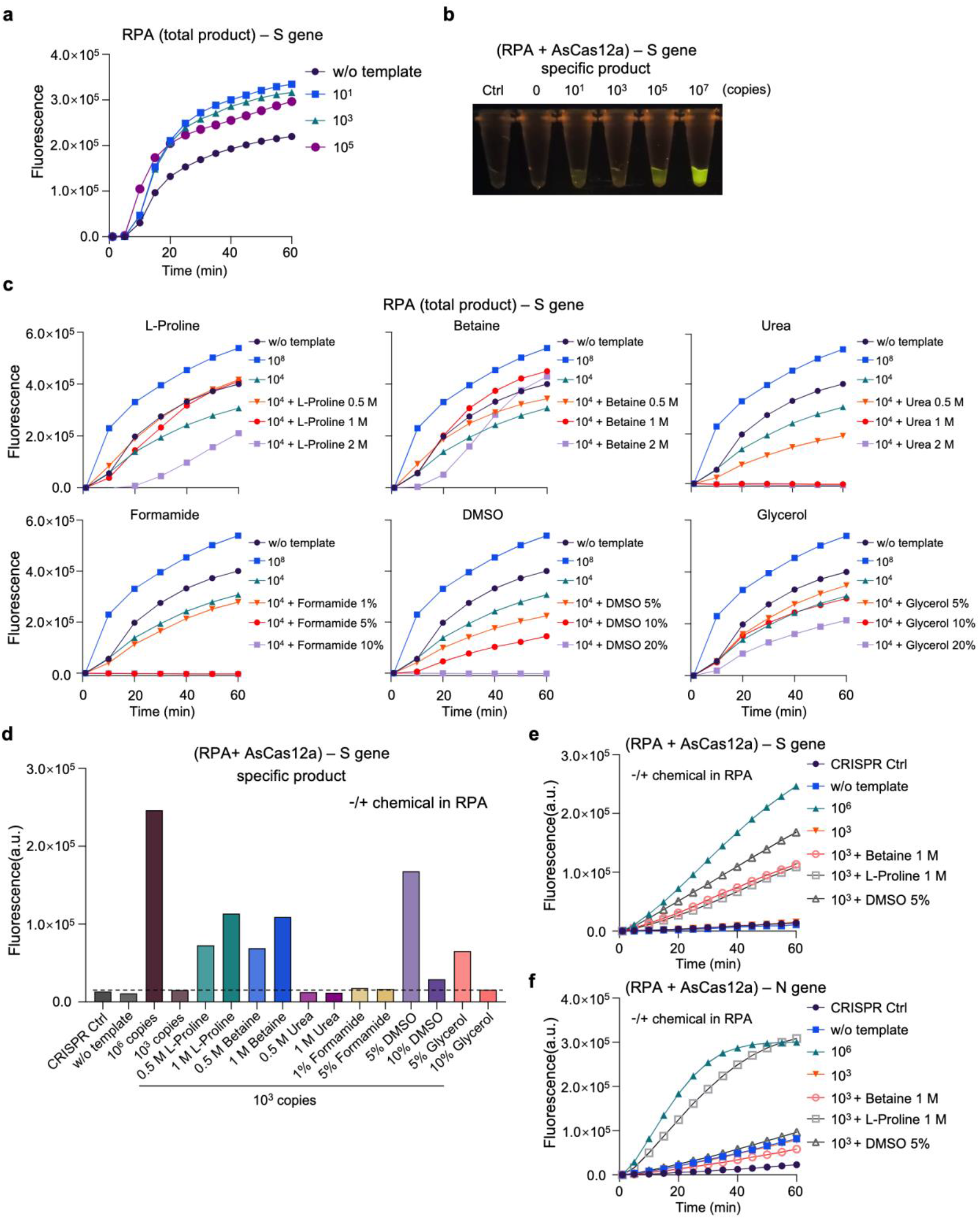
Evaluation of RPA and chemical additives for target detection. **a** Dynamic monitoring of total amplification product of RT-RPA by nonspecific DNA-binding dye EvaGreen using varied amount of SARS-CoV-2 S gene RNA template. **b** Specific target amplification is determined using purified RPA products by AsCas12a-based detection with fluorescence signal directly visualized by blue light illuminator. Varied amount of SARS-CoV-2 S gene RNA templates are used in RT-RPA for a reaction time of 30 min at 40°C before proceeding to purification and CRISPR detection. **c** Evaluation of chemical additives for the effects on total products of RT-RPA using SARS-CoV-2 S gene RNA template (10^4^ and 10^8^ copies). Indicated chemicals (+) are added in the RPA reaction mix and fluorescence kinetics is shown as determined by EvaGreen dye. w/o template means no template input in RPA reaction. **d** Evaluation of different chemical additives for the effect on specific target amplification as determined by AsCas12a-based CRISPR detection using purified PRA products (1:10 dilution) targeting SARS-CoV-2 S gene RNA. Indicated chemicals are included in the RPA reaction mix for a 30 min of reaction at 40°C. The RPA products are purified and then subjected to CRISPR detection. Endpoint (60 min) recording of fluorescence detection signals are shown. The dotted line indicates the fluorescence baseline of CRISPR detection signal from the sample using 10^3^ copies of SARS-CoV-2 synthetic RNA template. CRISPR Ctrl indicates no nucleic acid input for AsCas12a detection system; w/o template means no input for RPA reaction. **e** Fluorescence kinetics of AsCas12a-based detection is shown for the selected samples used in (d). **f** Fluorescence kinetics of AsCas12a-based detection are shown to detect specific target amplification from purified RT-RPA products using varied amount of SARS-CoV-2 N gene RNA template in the absence or presence of indicated chemicals. Indicated chemicals are included in the RPA reaction mix for a 30 min of reaction at 40°C. The RPA products are purified and then subjected to CRISPR detection.

**Supplementary Fig. 6.**
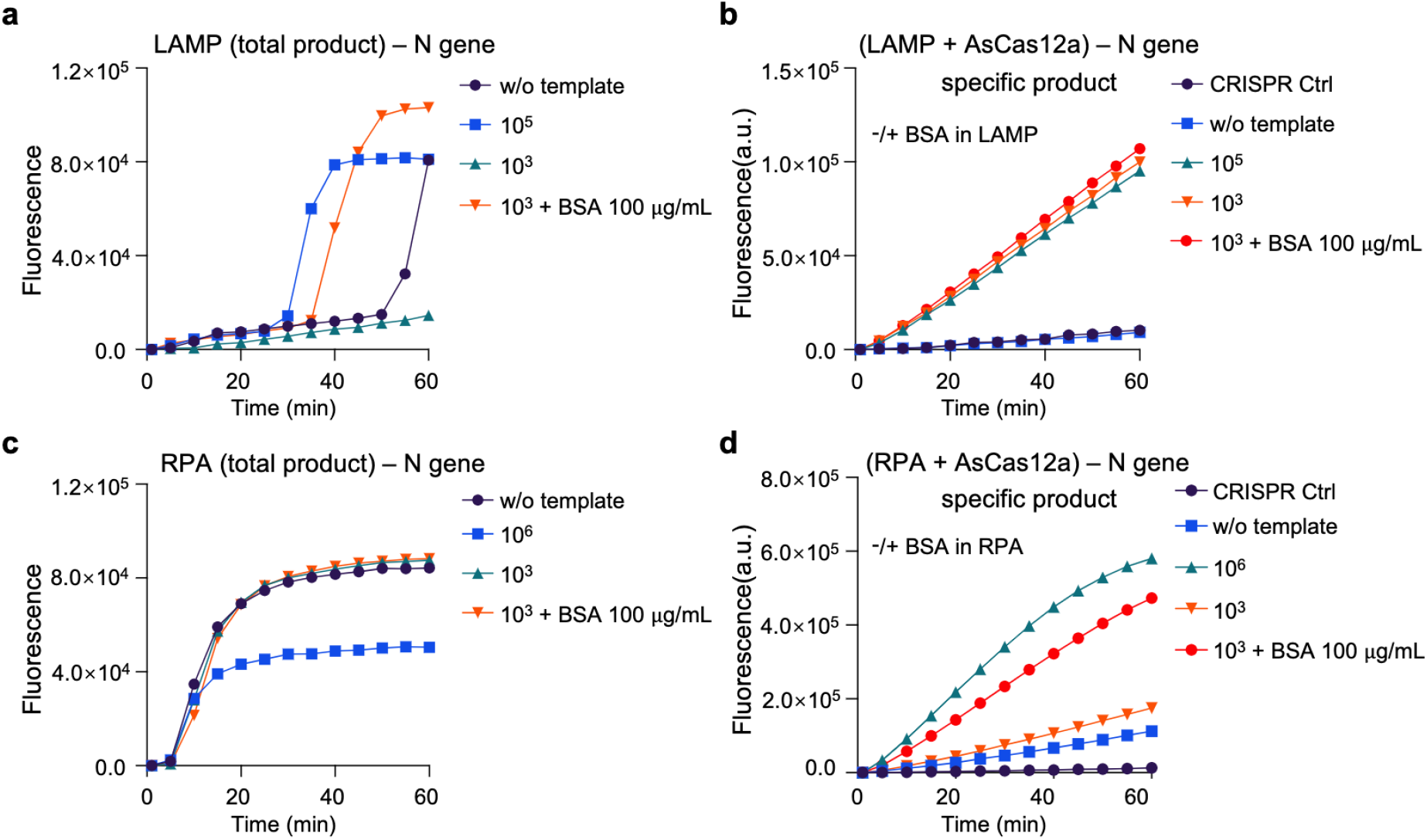
The effect of BSA on target amplification of LAMP and RPA. **a** Dynamic monitoring of total amplification product of RT-LAMP by nonspecific DNA-binding dye EvaGreen using varied amount of SARS-CoV-2 N gene RNA template in the absence or presence of BSA. w/o template means no template input in LAMP reaction. **b** Fluorescence kinetics of AsCas12a-based detection are shown to detect specific target amplification from purified RT-LAMP products (1:50 dilution) using varied amount of SARS-CoV-2 N gene RNA template. BSA is absent or present only in LAMP reaction mix. RT-LAMP reaction is performed for 15 min at 62°C before proceeding to purification and then CRISPR detection. CRISPR Ctrl indicates no nucleic acid input for AsCas12a detection system; w/o template means no input for LAMP reaction. **c** Dynamic monitoring of total amplification product of RT-RPA by nonspecific DNA-binding dye EvaGreen using varied amount of SARS-CoV-2 N gene RNA template in the absence or presence of BSA. w/o template means no template input in RPA reaction. **d** Fluorescence kinetics of AsCas12a-based detection are shown to detect specific target amplification from purified RT-RPA products using varied amount of SARS-CoV-2 N gene RNA template. BSA is absent or present only in RPA reaction mix. RT-RPA reaction is performed for 30 min at 40°C before proceeding to purification and then CRISPR detection. CRISPR Ctrl indicates no nucleic acid input for AsCas12a detection system; w/o template means no input for RPA reaction.

**Supplementary Fig. 7.**
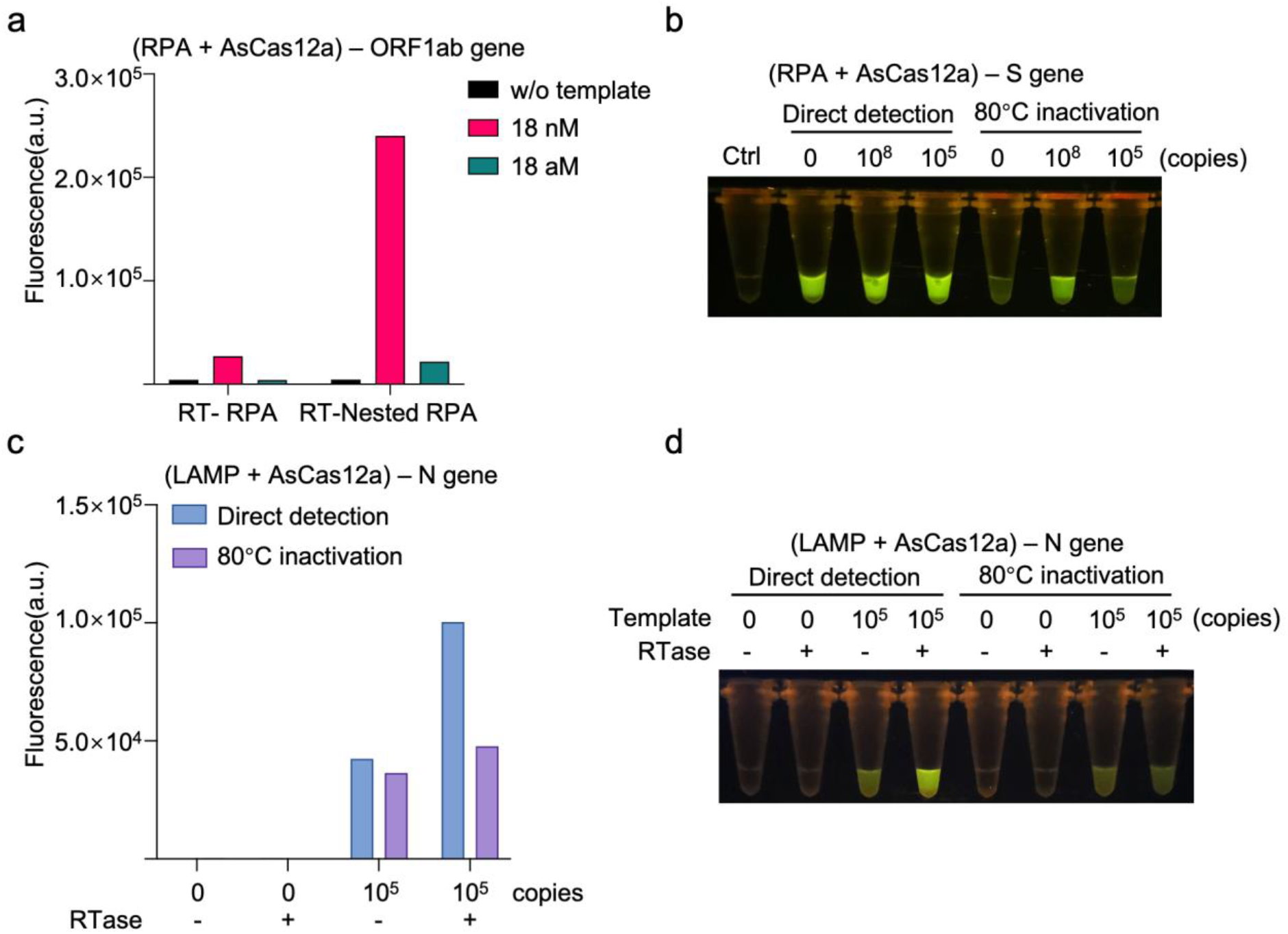
Performance of RT-Nested RPA and background removal for CRISPR detection. **a** Comparison of RT-RPA and RT-Nested RPA with varied amount of SARS-CoV-2 ORF1ab gene RNA template. RPA reaction is performed for 10 min at 37°C, and nested RPA has an additional round of pre-amplification for 5 min. 1 μL amplification product is subjected to AsCas12a-based detection and the endpoint (2h) fluorescence signals are shown. **b** Background signal removal by 80°C heat-inactivation for RPA-AsCas12a two-step detection assays to examine SARS-CoV-2 S gene RNA template. RT-RPA is performed at 40°C for 30 min and 2 μL amplification product is then subjected to AsCas12a-mediated CRISPR detection. Blue light illuminator is used to visualize the endpoint (60 min) fluorescence signal. **c, d** The effect of reverse transcriptase (RTase) on the background signal of LAMP-AsCas12a-based two-step CRISPR detection system. SARS-CoV-2 N gene DNA template is used for LAMP. The LAMP reaction is performed at 62°C for 15 min and 2 μL amplification product is then subjected to AsCas12a-mediated CRISPR detection. RTase is absent (-) or present (+) only in the CRISPR detection mix. The endpoint (60 min) fluorescence signals are shown by either bar plot (c) or direct visualization under blue light illuminator (d).

**Supplementary Table 1.** The constitutions of different reaction buffers.

| Buffer composition | NEBuffer 2 | NEBuffer 2.1 | NEBuffer 3 | NEBuffer 3.1 | Takara T | Takara K | Isothermal buffer | CutSmart |
| --- | --- | --- | --- | --- | --- | --- | --- | --- |
| Magnesium Acetate |  |  |  |  | 10 mM |  |  | 10 mM |
| MgCl<sub>2</sub> | 10 mM | 10 mM | 10 mM | 10 mM |  | 10 mM |  |  |
| Potassium Acetate |  |  |  |  | 66 mM |  |  | 50 mM |
| KCl |  |  |  |  |  | 100 mM | 50 mM |  |
| NaCl | 50 mM | 50 mM | 100 mM | 100 mM |  |  |  |  |
| Tris-acetate |  |  |  |  | 33 mM |  |  | 20 mM |
| Tris-HCl | 10 mM | 10 mM | 50 mM | 50 mM |  | 20 mM | 20 mM |  |
| (NH<sub>4</sub>)<sub>2</sub>SO<sub>4</sub> |  |  |  |  |  |  | 10 mM |  |
| MgSO<sub>4</sub> |  |  |  |  |  |  | 2 mM |  |
| Tween 20 |  |  |  |  |  |  | 0.1% |  |
| DTT | 1 mM |  | 1 mM |  | 0.5 mM | 1 mM |  |  |
| BSA |  | 100 µg/mL |  | 100 µg/mL |  |  |  | 100 µg/mL |
| pH | 7.9 | 7.9 | 7.9 | 7.9 | 7.9 | 8.5 | 8.8 | 7.9 |

**Supplementary Table 2.**
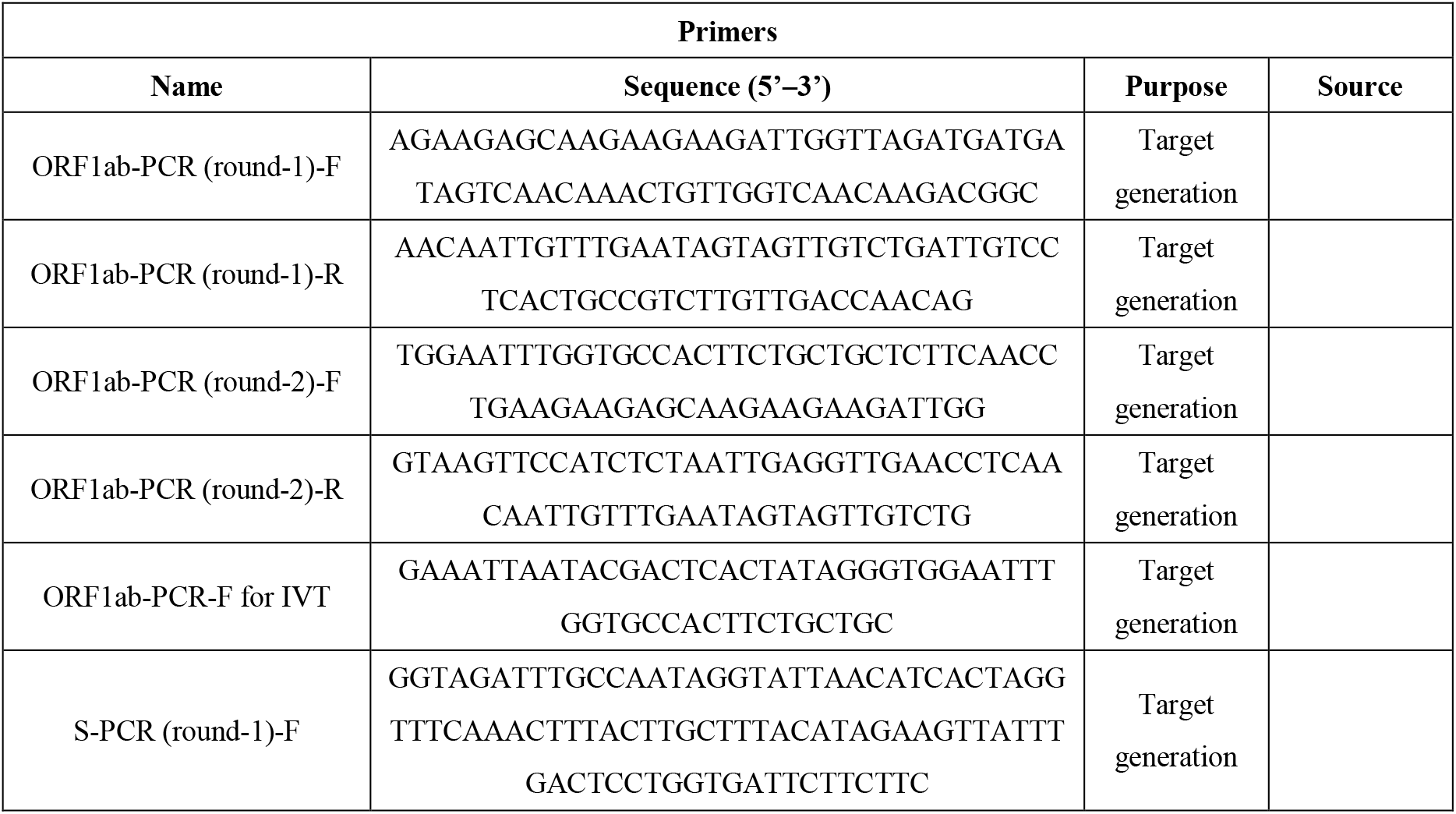
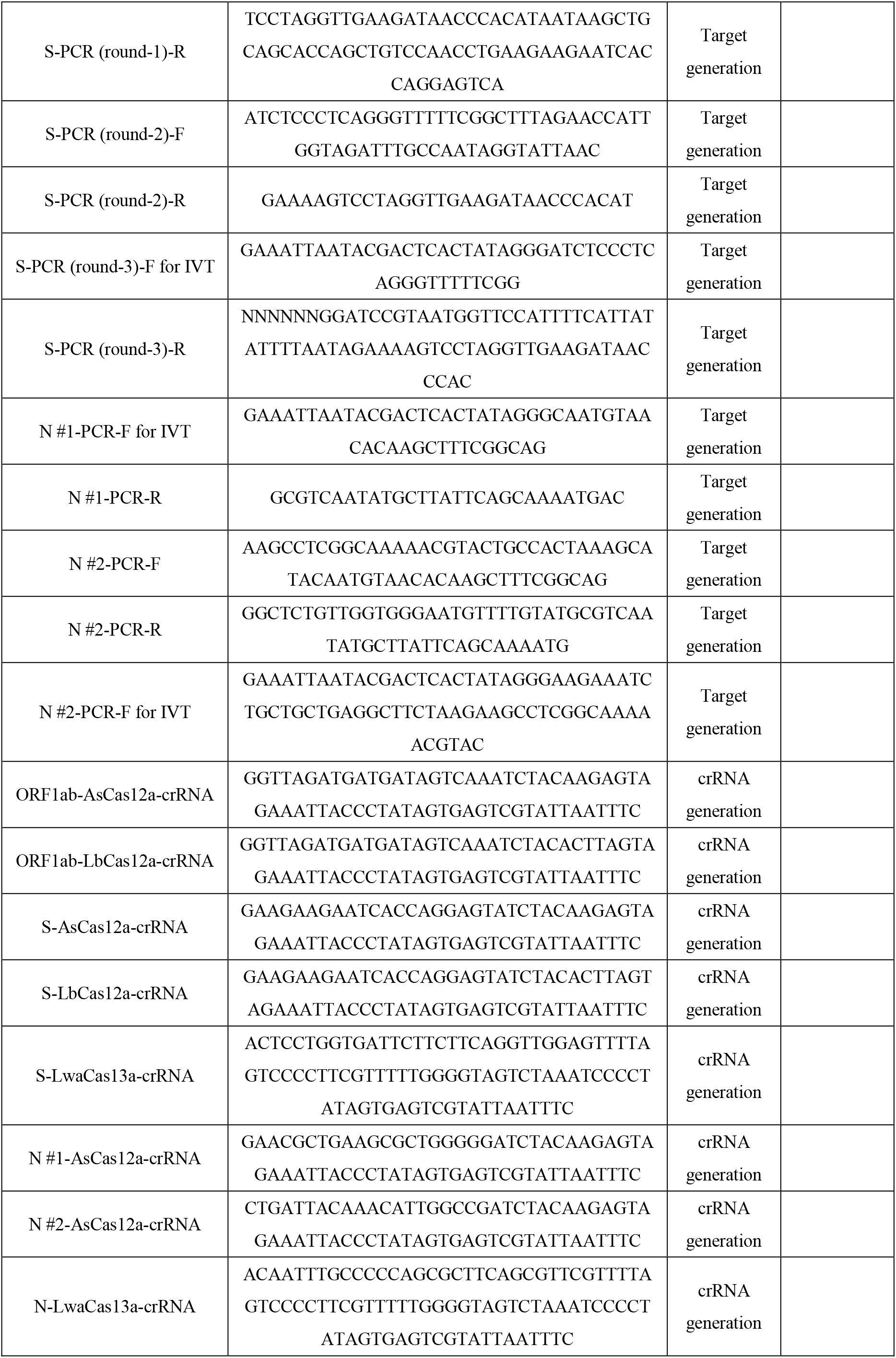
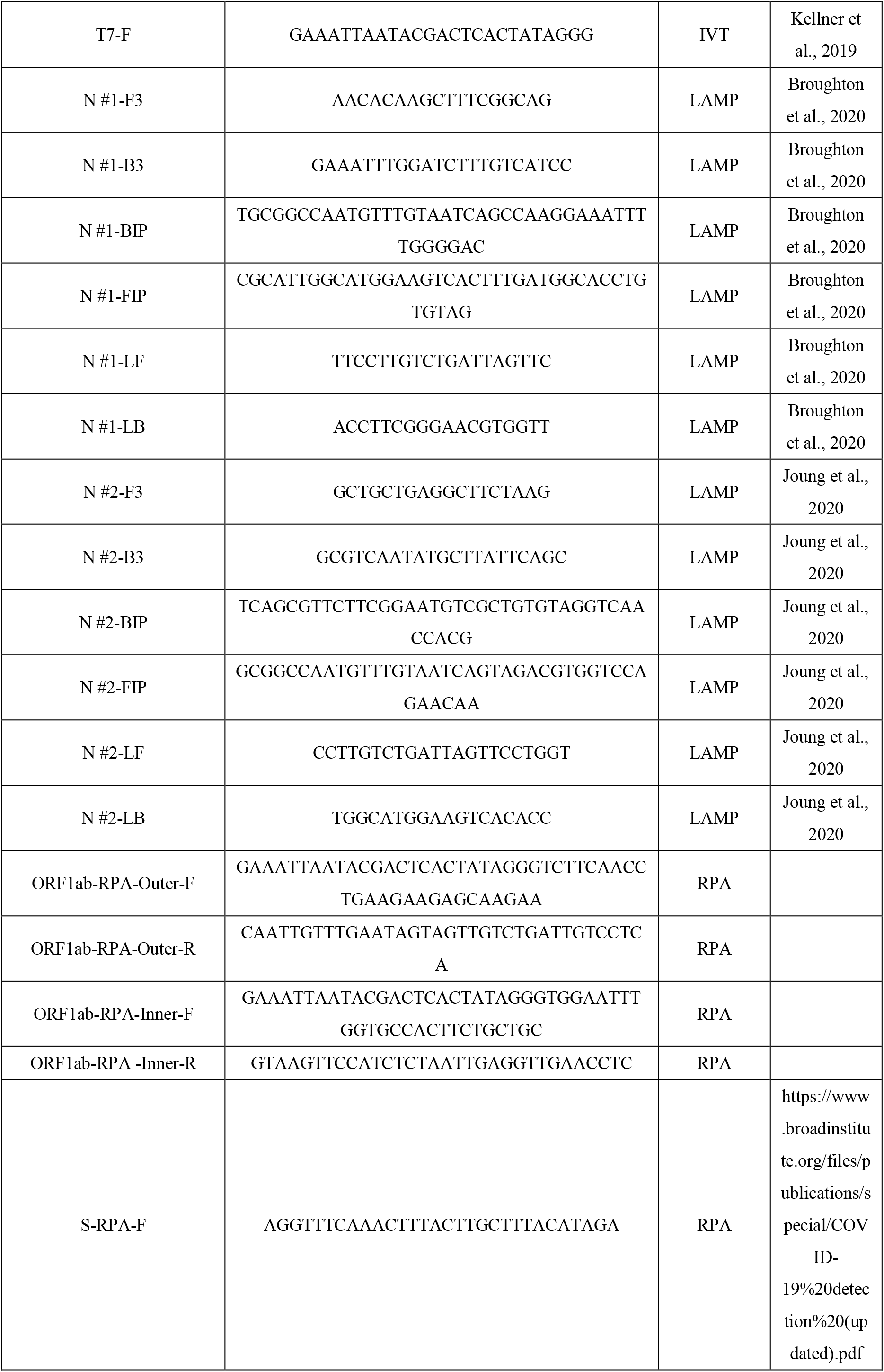
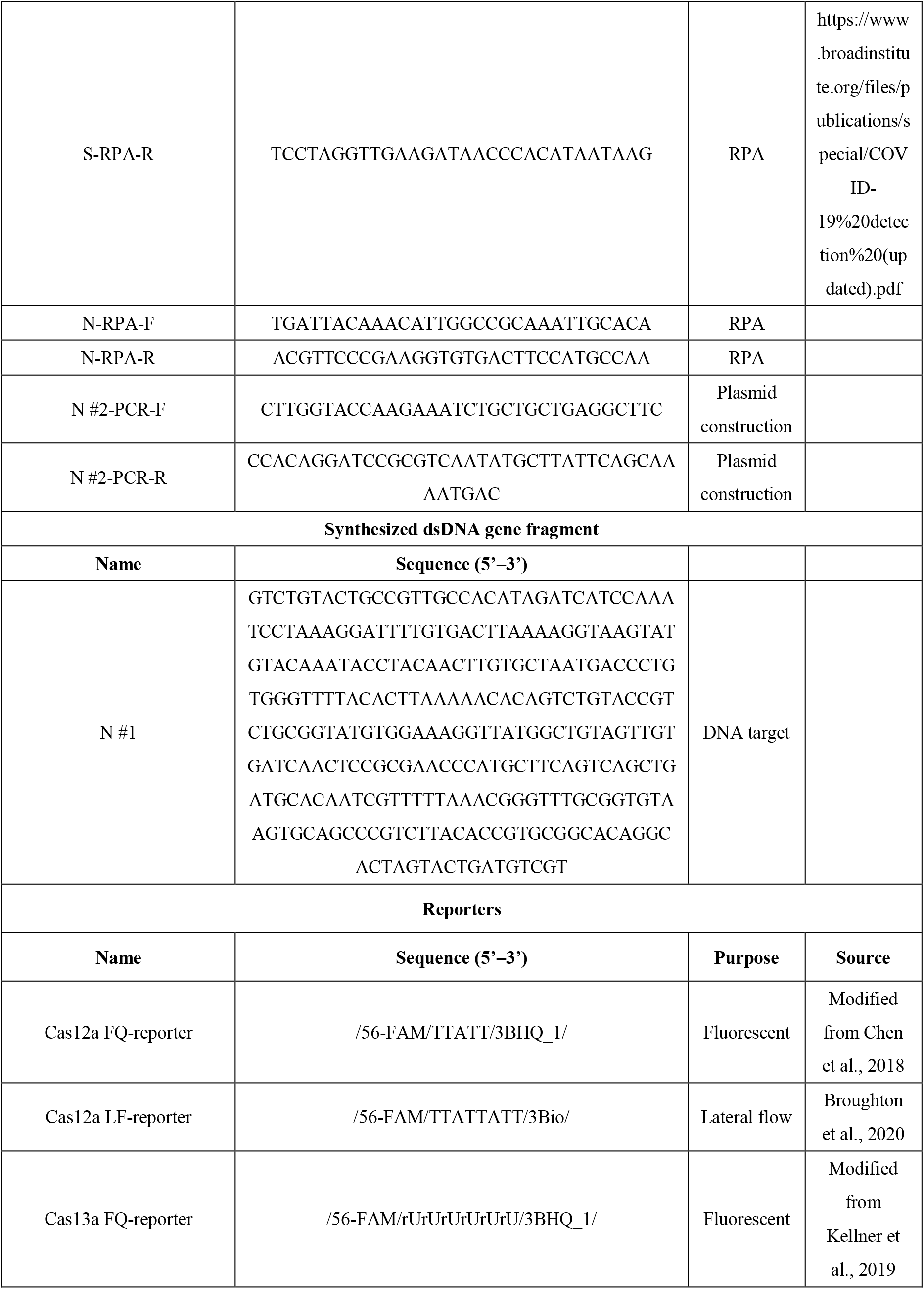
Oligonucleotides used in this study.

**Supplementary Table 3.** Reagents and instruments used in this study.

| Reagent/Instrument name | Commercial vendor | Item No. |
| --- | --- | --- |
| 10x T buffer | Takara Bio | SD6092 |
| 10x K buffer | Takara Bio | SD0008 |
| 10x Isothermal Amplification Buffer | New England Biolab® Inc | B0374S |
| CutSmart® Buffer | New England Biolab® Inc | B7204S |
| NEBuffer™ 2 | New England Biolab® Inc | B7002S |
| NEBuffer™ 2.1 | New England Biolab® Inc | B7202S |
| NEBuffer™ 3 | New England Biolab® Inc | B7003S |
| NEBuffer™ 3.1 | New England Biolab® Inc | B7203S |
| Bst 2.0 WarmStart DNA Polymerase | New England Biolab® Inc | M0538S |
| Phanta Max Super-Fidelity DNA Polymerase | Vazyme | P505 |
| WarmStart RTx Reverse Transcriptase | New England Biolab® Inc | M0380S |
| NxGen® T7 RNA polymerase | Lucigen | 30223-1 |
| ProtoScript II Reverse Transcriptase | New England Biolab® Inc | M0368S |
| High-Capacity cDNA Reverse Transcription Kit | Thermo Fisher | 4368813 |
| T4 DNA ligase | Thermo Fisher | EL0016 |
| RNaseOUT™ Recombinant Ribonuclease Inhibitor | Thermo Fisher | 10777019 |
| BamH I restriction enzyme | Takara Bio | 1010A |
| Kpn I restriction enzyme | Takara Bio | 1068A |
| Xho I restriction enzyme | Takara Bio | 1094B |
| Rosetta 2(DE3)pLySs | Youbio | ST1147 |
| Isopropyl β-D-thiogalactoside | Solarbio | I8070 |
| Lysozyme | Solarbio | L8120 |
| PMSF | Beyotime | ST505 |
| BCA protein assay kit with BSA | Meilunbio | MA0082 |
| HisPur™ Cobalt Resin | Thermo Fisher | 89964 |
| Zeba™ Spin Desalting Columns, 7K MWCO, 2 mL | Thermo Fisher | 89889 |
| Amicon® Ultra-15 Centrifugal Filter Unit | Milipore | UFC905008 |
| Lipofectamine™ 2000 Transfection Reagent | Thermo Fisher | 11668019 |
| Opti-MEM™, Reduced Serum Medium, no phenol red | Thermo Fisher | 11058021 |
| L-Proline | Solarbio | SP8540 |
| Betaine | Solarbio | B8230 |
| Urea | Solarbio | U8020 |
| Formamide | Sigma Aldrich | F9037 |
| Dimethyl sulfoxide | Solarbio | D8370 |
| Glycerol | Sangon Biotech | A600232 |
| Bovine serum albumin | New England Biolab® Inc | B9000S |
| qPCR Lentivirus Titer Kit | Applied Biological Materials Inc | LV900 |
| RNA Clean & Concentrator™-5 | ZYMO RESEARCH | R1013 |
| GeneJET PCR Purification Kit | Thermo Fisher | K0702 |
| TwistAmp® Basic | TwistDx™ | tabas03kit |
| EvaGreen® Dye, 20x in Water | Biotium | 31000 |
| HybriDetect-Universal Lateral Flow Assay Kit | Milenia Biotec GmbH | MGHD 1 |
| UltraPure™ DNase/RNase-Free Distilled Water | Thermo Fisher | 10977015 |
| E-Gel™ Safe Imager™ Real-Time Transilluminator | Thermo Fisher | G6500 |
| Qubit® 3.0 Fluorometer | Thermo Fisher | Q33216 |
| QuantStudio™ 5 Real-Time PCR System | Thermo Fisher | A28139 |
| Ultrasonic cell disruptor | Ningbo Scientz Biotechnology | II D |

## References

1. Shin DJ, Andini N, Hsieh K, Yang S, Wang TH. Emerging Analytical Techniques for Rapid Pathogen Identification and Susceptibility Testing. Annual review of analytical chemistry 12, 41–67 (2019).

2. Chen H, Liu K, Li Z, Wang P. Point of care testing for infectious diseases. Clinica chimica acta; international journal of clinical chemistry 493, 138–147 (2019).

3. Vandenberg O, Martiny D, Rochas O, van Belkum A, Kozlakidis Z. Considerations for diagnostic COVID-19 tests. Nature reviews Microbiology 19, 171–183 (2021).

4. Wang X, Shang X, Huang X. Next-generation pathogen diagnosis with CRISPR/Cas-based detection methods. Emerging microbes & infections 9, 1682–1691 (2020).

5. Teng F, et al. CDetection: CRISPR-Cas12b-based DNA detection with sub-attomolar sensitivity and single-base specificity. Genome biology 20, 132 (2019).

6. Chen JS, et al. CRISPR-Cas12a target binding unleashes indiscriminate single-stranded DNase activity. Science 360, 436–439 (2018).

7. Li SY, et al. CRISPR-Cas12a-assisted nucleic acid detection. Cell discovery 4, 20 (2018).

8. Li L, et al. HOLMESv2: A CRISPR-Cas12b-Assisted Platform for Nucleic Acid Detection and DNA Methylation Quantitation. ACS synthetic biology 8, 2228–2237 (2019).

9. Gootenberg JS, Abudayyeh OO, Kellner MJ, Joung J, Collins JJ, Zhang F. Multiplexed and portable nucleic acid detection platform with Cas13, Cas12a, and Csm6. Science 360, 439–444 (2018).

10. Gootenberg JS, et al. Nucleic acid detection with CRISPR-Cas13a/C2c2. Science 356, 438–442 (2017).

11. Harrington LB, et al. Programmed DNA destruction by miniature CRISPR-Cas14 enzymes. Science 362, 839–842 (2018).

12. Kellner MJ, Koob JG, Gootenberg JS, Abudayyeh OO, Zhang F. SHERLOCK: nucleic acid detection with CRISPR nucleases. Nature protocols 14, 2986–3012 (2019).

13. Wang M, Zhang R, Li J. CRISPR/cas systems redefine nucleic acid detection: Principles and methods. Biosensors & bioelectronics 165, 112430 (2020).

14. van Dongen JE, Berendsen JTW, Steenbergen RDM, Wolthuis RMF, Eijkel JCT, Segerink LI. Point-of-care CRISPR/Cas nucleic acid detection: Recent advances, challenges and opportunities. Biosensors & bioelectronics 166, 112445 (2020).

15. Broughton JP, et al. CRISPR-Cas12-based detection of SARS-CoV-2. Nature biotechnology 38, 870–874 (2020).

16. Crone MA, et al. A role for Biofoundries in rapid development and validation of automated SARS-CoV-2 clinical diagnostics. Nature communications 11, 4464 (2020).

17. Patchsung M, et al. Clinical validation of a Cas13-based assay for the detection of SARS-CoV-2 RNA. Nature biomedical engineering 4, 1140–1149 (2020).

18. Nguyen LT, Smith BM, Jain PK. Enhancement of trans-cleavage activity of Cas12a with engineered crRNA enables amplified nucleic acid detection. Nature communications 11, 4906 (2020).

19. Guo L, et al. SARS-CoV-2 detection with CRISPR diagnostics. Cell discovery 6, 34 (2020).

20. Ding X, et al. Ultrasensitive and visual detection of SARS-CoV-2 using all-in-one dual CRISPR-Cas12a assay. Nature communications 11, 4711 (2020).

21. Arizti-Sanz J, et al. Streamlined inactivation, amplification, and Cas13-based detection of SARS-CoV-2. Nature communications 11, 5921 (2020).

22. Wang R, et al. opvCRISPR: One-pot visual RT-LAMP-CRISPR platform for SARS-cov-2 detection. Biosensors & bioelectronics 172, 112766 (2021).

23. Fozouni P, et al. Amplification-free detection of SARS-CoV-2 with CRISPR-Cas13a and mobile phone microscopy. Cell 184, 323–333 e329 (2021).

24. Ramachandran A, et al. Electric field-driven microfluidics for rapid CRISPR-based diagnostics and its application to detection of SARS-CoV-2. Proceedings of the National Academy of Sciences of the United States of America 117, 29518–29525 (2020).

25. Joung J, et al. Detection of SARS-CoV-2 with SHERLOCK One-Pot Testing. The New England journal of medicine 383, 1492–1494 (2020).

26. Ma P, et al. MeCas12a, a Highly Sensitive and Specific System for COVID-19 Detection. Advanced science, 2001300 (2020).

27. Simpson RJ. Stabilization of proteins for storage. Cold Spring Harbor protocols 2010, pdb top79 (2010).

28. Varadharajan B, Parani M. DMSO and betaine significantly enhance the PCR amplification of ITS2 DNA barcodes from plants. Genome 64, 165–171 (2021).

29. Jensen MA, Fukushima M, Davis RW. DMSO and betaine greatly improve amplification of GC-rich constructs in de novo synthesis. PloS one 5, e11024 (2010).

30. Ralser M, Querfurth R, Warnatz HJ, Lehrach H, Yaspo ML, Krobitsch S. An efficient and economic enhancer mix for PCR. Biochemical and biophysical research communications 347, 747–751 (2006).

31. Henke W, Herdel K, Jung K, Schnorr D, Loening SA. Betaine improves the PCR amplification of GC-rich DNA sequences. Nucleic acids research 25, 3957–3958 (1997).

32. Nagai M, Yoshida A, Sato N. Additive effects of bovine serum albumin, dithiothreitol, and glycerol on PCR. Biochemistry and molecular biology international 44, 157–163 (1998).

33. Luo GC, Yi TT, Jiang B, Guo XL, Zhang GY. Betaine-assisted recombinase polymerase assay with enhanced specificity. Analytical biochemistry 575, 36–39 (2019).

34. Liu KI, et al. A chemical-inducible CRISPR-Cas9 system for rapid control of genome editing. Nature chemical biology 12, 980–987 (2016).

35. Maji B, et al. Multidimensional chemical control of CRISPR-Cas9. Nature chemical biology 13, 9–11 (2017).

36. Yin H, et al. Structure-guided chemical modification of guide RNA enables potent non-viral in vivo genome editing. Nature biotechnology 35, 1179–1187 (2017).

37. Zhang S, Shen J, Li D, Cheng Y. Strategies in the delivery of Cas9 ribonucleoprotein for CRISPR/Cas9 genome editing. Theranostics 11, 614–648 (2021).

38. Ma C, Wang Y, Zhang P, Shi C. Accelerated isothermal nucleic acid amplification in betaine-free reaction. Analytical biochemistry 530, 1–4 (2017).

39. Farell EM, Alexandre G. Bovine serum albumin further enhances the effects of organic solvents on increased yield of polymerase chain reaction of GC-rich templates. BMC research notes 5, 257 (2012).

40. Kageyama K, Komatsu T, Suga H. Refined PCR protocol for detection of plant pathogens in soil. Journal of General Plant Pathology 69, 153–160 (2003).

41. Plante D, Belanger G, Leblanc D, Ward P, Houde A, Trottier YL. The use of bovine serum albumin to improve the RT-qPCR detection of foodborne viruses rinsed from vegetable surfaces. Letters in applied microbiology 52, 239–244 (2011).

42. Tarelli E, et al. Recombinant human albumin as a stabilizer for biological materials and for the preparation of international reference reagents. Biologicals: journal of the International Association of Biological Standardization 26, 331–346 (1998).

43. Marth E, Kleinhappl B. Albumin is a necessary stabilizer of TBE-vaccine to avoid fever in children after vaccination. Vaccine 20, 532–537 (2001).

44. Morita Y, Nakamori S, Takagi H. L-proline accumulation and freeze tolerance of Saccharomyces cerevisiae are caused by a mutation in the PRO1 gene encoding gamma-glutamyl kinase. Applied and environmental microbiology 69, 212–219 (2003).

45. Zosel F, Mercadante D, Nettels D, Schuler B. A proline switch explains kinetic heterogeneity in a coupled folding and binding reaction. Nature communications 9, 3332 (2018).

46. Ben-Gedalya T, et al. Alzheimer’s disease-causing proline substitutions lead to presenilin 1 aggregation and malfunction. The EMBO journal 34, 2820–2839 (2015).

47. Gladkevich A, Bosker F, Korf J, Yenkoyan K, Vahradyan H, Aghajanov M. Proline-rich polypeptides in Alzheimer’s disease and neurodegenerative disorders -- therapeutic potential or a mirage? Progress in neuro-psychopharmacology & biological psychiatry 31, 1347–1355 (2007).

48. Bolli R, Woodtli K, Bartschi M, Hofferer L, Lerch P. L-Proline reduces IgG dimer content and enhances the stability of intravenous immunoglobulin (IVIG) solutions. Biologicals: journal of the International Association of Biological Standardization 38, 150–157 (2010).

49. Berger M. L-proline-stabilized human IgG: Privigen(R) 10% for intravenous use and Hizentra(R) 20% for subcutaneous use. Immunotherapy 3, 163–176 (2011).

50. Hagan JB, et al. Safety of L-proline as a stabilizer for immunoglobulin products. Expert review of clinical immunology 8, 169–178 (2012).

51. Rao SN, Mohan DC, Adimurthy S. L-proline: an efficient catalyst for transamidation of carboxamides with amines. Organic letters 15, 1496–1499 (2013).

52. Yang JW, Chandler C, Stadler M, Kampen D, List B. Proline-catalysed Mannich reactions of acetaldehyde. Nature 452, 453–455 (2008).

53. Tanimoro K, Ueno M, Takeda K, Kirihata M, Tanimori S. Proline catalyzes direct C-H arylations of unactivated arenes. The Journal of organic chemistry 77, 7844–7849 (2012).

54. Dasgupta M, Kishore N. Selective inhibition of aggregation/fibrillation of bovine serum albumin by osmolytes: Mechanistic and energetics insights. PloS one 12, e0172208 (2017).

55. Kumar TK, Samuel D, Jayaraman G, Srimathi T, Yu C. The role of proline in the prevention of aggregation during protein folding in vitro. Biochemistry and molecular biology international 46, 509–517 (1998).

56. Abudayyeh OO, et al. RNA targeting with CRISPR-Cas13. Nature 550, 280–284 (2017).

57. Li SY, Cheng QX, Liu JK, Nie XQ, Zhao GP, Wang J. CRISPR-Cas12a has both cis- and trans-cleavage activities on single-stranded DNA. Cell research 28, 491–493 (2018).

